# Symbiosis, Selection and Novelty: Freshwater Adaptation in the Unique Sponges of Lake Baikal

**DOI:** 10.1101/416230

**Authors:** Nathan J Kenny, Bruna Plese, Ana Riesgo, Valeria B. Itskovich

## Abstract

Freshwater sponges (Spongillida) are a unique lineage of demosponges that secondarily colonized lakes and rivers and are now found ubiquitously in these ecosystems. They developed specific adaptations to freshwater systems, including the ability to survive extreme thermal ranges, long-lasting dessication, anoxia, and resistance to a variety of pollutants. While spongillids have colonized all freshwater systems, the family Lubomirskiidae is endemic to Lake Baikal, and plays a range of key roles in this ecosystem. Our work compares the genomic content and microbiome of individuals of three species of the Lubomirskiidae, providing hypotheses for how molecular evolution has allowed them to adapt to their unique environments. We have sequenced deep (>92% of the metazoan ‘Benchmarking Universal Single-Copy Orthologs’ (BUSCO) set) transcriptomes from three species of Lubomirskiidae and a draft genome resource for *Lubomirskia baikalensis*. We note Baikal sponges contain unicellular algal and bacterial symbionts, as well as the dinoflagellate *Gyrodinium*. We investigated molecular evolution, gene duplication and novelty in freshwater sponges compared to marine lineages. Sixty one orthogroups have consilient evidence of positive selection. Transporters (e.g. *zinc transporter-2),* transcription factors (*aristaless-related homeobox*) and structural proteins (for example *actin-3*), alongside other genes, are under strong evolutionary pressure in freshwater, with duplication driving novelty across the Spongillida, but especially in the Lubomirskiidae. This addition to knowledge of freshwater sponge genetics provides a range of tools for understanding the molecular biology and, in the future, the ecology (for example, colonization and migration patterns) of these key species.

## Introduction

Freshwater sponges belong to the monophyletic order Spongillida, a unique lineage of demosponges that colonized lentic and lotic systems sometime in the Permo-Carboniferous (around 311 million years ago; Schuster et al. 2018). To be able survive such a drastic change in environment, freshwater sponges adapted at the molecular, physiological, and structural level. They are able to survive a wide range of thermal conditions, including permafrost, fluctuating water levels, anoxia, and certain levels of chemicals and pollutants in the water (Manconi and Pronzato, 2008). Among their physical adaptations, freshwater sponges have developed gemmules to enable them to endure winter conditions, encapsulating undifferentiated sponge cells into silica structures (Manconi and Pronzato, 2008). The sponge family Lubomirskiidae (Demospongiae: Spongillida) is endemic to Lake Baikal, southern Siberia, and is a vital part of that unusual ecosystem. They colonized this lake approximately 3.4 MYA, approximately 15 million years after the recent radiation of freshwater sponges (Schuster et al 2018). This family contains four genera and thirteen currently accepted species (Itskovich et al 2015). As sponges are the most prominent component of the benthic assemblage of Lake Baikal, making up around 44% of its biomass (Pile et al 1997), they play a vital role within this ecological community.

The Lubomirskiidae possess a number of unique features. They are large for freshwater sponges, growing to around 1 meter in size (Kozhov 1963). They, as with all freshwater sponges, depend on symbiotic associations with bacteria and various chlorophyll-producing microorganisms: green algae, dinoflagellates and diatoms (Pile et al 1997) and their diversification is yet to be fully understood. They can be described as a “species flock” (Schröder et al 2003), and recent studies have found discrepancies between molecular and morphological data used to define species boundaries in Baikalian sponges (Itskovich et al 2015). Among their morphological adaptations is the loss of the ability to develop gemmules (Pronzato and Manconi 2008).

*Lubomirskia baikalensis* (Fig 1A) is the most abundant and best studied species of the Lubomirskiidae. It is found throughout the lake, but its morphology varies according to depth (Kozhov 1963). It grows as a mat in shallower waters, while in deeper areas of the lake it increases in height and begins to branch. It can form dense aggregates in favourable conditions, and forms beds providing the habitat for a range of other species. Like other members of the Lubomirskiidae, *L. baikalensis* possesses symbiotic dinoflagellates assumed to be vital for proper biological function (Annenkova et al 2011, Chernogor et al 2013). Symbiotic bacteria also contribute to the success of this clade, as they do in other members of the Spongillidae (e.g. Latyshev et al 1992, Gernert et al 2005, Kaluzhnaya et al 2011, 2012, Costa et al 2013 Kaluzhnaya and Itskovich 2015).

**Figure 1:**
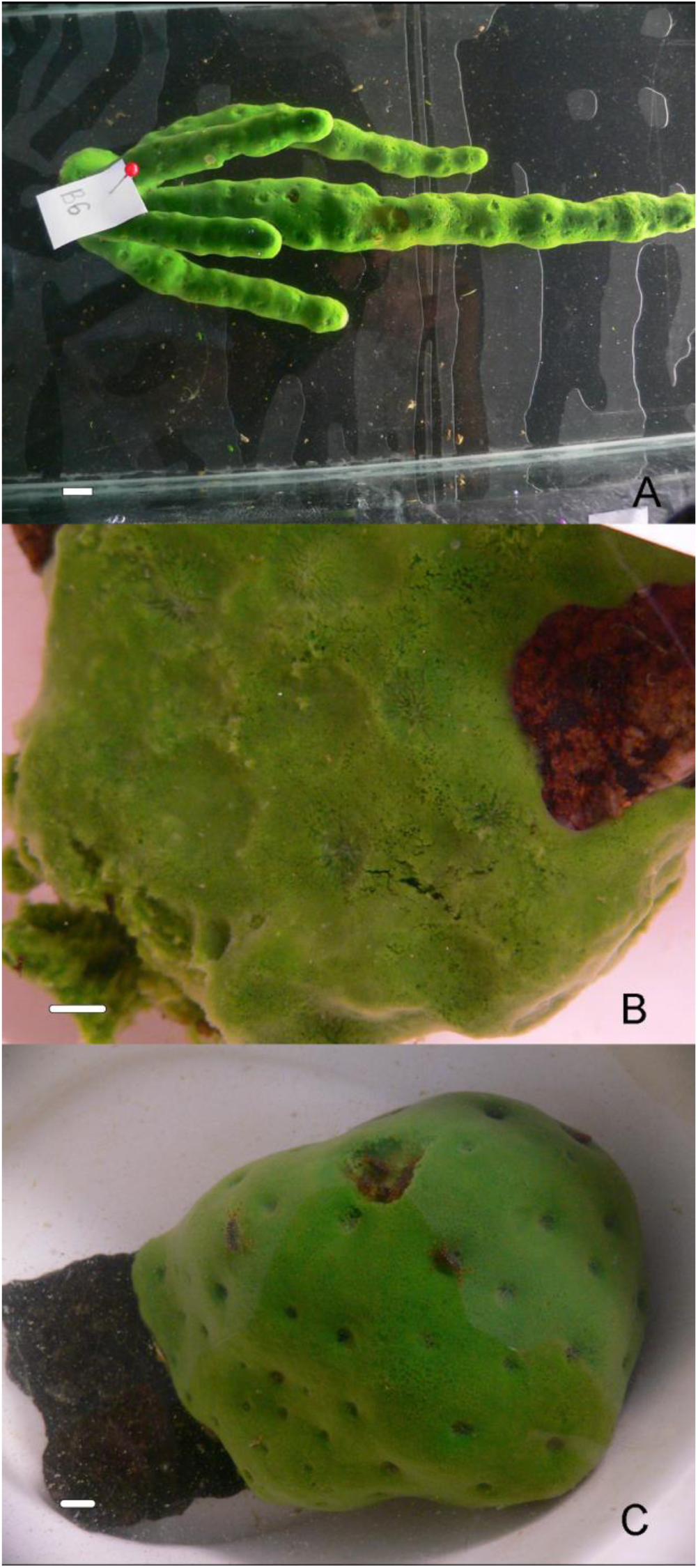
Specimens used for RNA and gDNA extraction. A) *Lubomirskia baikalensis*. B) *Lubomirskia abietina*. C) *Baikalospongia bacillifera*. Scale bar lengths 1 cm.

Lake Baikal is the largest freshwater lake in the world by volume, the deepest and the oldest (Timoshkin, 2001, Rusinek et al., 2012a). This has allowed a large number of species to adapt to its unique conditions (Timoshkin, 2001; Rusinek et al., 2012b). The lake has exhibited approximately the same conditions for the past 2 – 4 million years (Kozhova and Izmest’eva, 1998), separated from other major freshwater ecosystems (Timofeyev, 2010). The lake itself freezes over in winter (Kozhov 1963) but regions deeper than 250m are a constant 3.3-4.3°C (Shimaraev et al., 1994). Summer temperatures can reach as high as 20° C in small bays (Pomazkina et al 2012), although a maximum of 17° C is more common (Kozhova and Izmest’eva, 1998). Coupled with high oxygen levels throughout the water column and generally oligotrophic conditions (Kozhov, 1963), these circumstances provide a challenging environment. In the last century, the Lake Baikal ecosystem has been subject to a variety of negative environmental influences. This includes pollution from both agriculture and industry, varying water levels as a result of damming and agricultural irrigation, eutrophication as a result of fertilizer run off, and algal blooms, including those by invasive species (for example, Romanova et al 2015, Ciesielski et al 2016, Kasimov et al 2017). Sponges, with their filter-feeding lifestyle, and particularly the Lubomirskiidae, with their reliance on symbionts, may be particularly vulnerable to such pollution. Bleaching in particular has been a major issue for the Lubomirskiidae in recent decades (Kaluzhnaya and Itskovich 2015, Khanaev et al 2017). Cases of novel diseases have also been reported (e.g. Denikina et al 2016, Kaluzhnaya and Itskovich 2015, Kaluzhnaya and Itskovich 2017, Kulakova et al 2017, Itskovich et al 2018). These changes are often associated with widespread changes to the presence and the ratio of sponge-associated bacterial communities (Denikina et al 2016, Kaluzhnaya and Itskovich 2015, Kulakova et al 2017). It is vital to understand the speciation patterns and diversity of the Lubomirskiidae in order to understand the impact of disease, manage conservation efforts and ensure the survival of the Baikal ecosystem (Khanaev et al 2017).

The biology of Baikal-endemic species has been studied from a variety of angles in the past decades (e.g. Rusinek et al., 2012b). However, only in the last few years has it been possible to study the molecular adaptations of these species, as the rise of “omic” technologies has allowed the study of even recalcitrant and rare species. The affordability of these and associated computational improvements has made such an approach attractive (Goodwin et al 2016). This has lead to the investigation of a number of Baikal species using next-generation sequencing (e.g. Rivarola-Duarte et al 2014, Romanova et al 2016, Naumenko et al 2017).

Such technologies have been highly successful in investigating poriferan diversity (e.g. Riesgo et al 2014, Fernandez-Valverde et al 2015, Kenny et al 2017 and others). However, to date, there have been no studies on sponges from Lake Baikal, and no freshwater sponge genomes are presently extant in the published record.

To increase our understanding of the biology of these unique sponges from a range of perspectives, we have assembled and analysed transcriptomes of three Baikal-endemic species, *Baikalospongia bacillifera, Lubomirskia abietina* and *L. baikalensis* (Fig 1). Alongside these samples we have also assembled a draft genome for *L. baikalensis.* Our main objective was to reveal the specific molecular signatures of adaptation in freshwater sponges, and we have identified a number of key changes in genes involved in freshwater adaptation, in transcription factors, structural proteins, membrane transport molecules and other proteins. These sequences will help us understand the biology, symbioses, population dynamics and physiology of the idiosyncratic sponges of Lake Baikal, and freshwater sponges more generally.

## Results and Discussion

### Sequencing and Assembly Overview

Initial sequencing results were of variable quality, and required cleaning before use in our analyses. Full details regarding this process, and relevant statistics can be seen in Supplementary File 1. Both clean and original reads have been uploaded to the NCBI SRA, with accession number <u>PRJNA431612</u>. Metrics related to the assemblies of the transcriptomes studied in this work can be seen in Supplementary File 1, Table 2, and that file contains further details about assembly and analysis. Individual assemblies were made using Trinity for all transcriptome datasets, each representing a single sponge. Given that we wish to consider both bacterial and metazoan sequences in the present study, it is worth considering the relative GC% of reads and our assemblies. The GC% of bacterial genomes is often, but not always, higher than that of metazoans, and higher GC% can empirically be an indicator of higher levels of bacterial sequence in a sample. This is, however, not universal. The Flavobacteriaceae, for instance, commonly found in both marine and freshwater biomes, have GC% of 31-35% (e.g. Tekedar et al 2017).

The GC% of our paired transcriptomic reads remained stable at 49% for all samples both before and after read cleaning. This stability indicates that all samples were of comparable composition, and none contained any obvious levels of exogenous DNA or symbiotic content relative to the others. The GC% of our genomic reads declined slightly with read cleaning, from 43.5 to 41%. This finding is confirmed in our Blobplots (Fig 2), displaying assembled sequence data separated by coverage on one axis, and GC content on the other, allowing easy visualisation of content. This shows that the small amount of bacterial sequences present have a very similar GC% to our assembled reads. This prohibits us from “binning” bacterial sequences by GC% to construct separate assemblies, but bacterial sequences could be recognised by BLAST similarity, as discussed elsewhere in this work.

**Figure 2:**
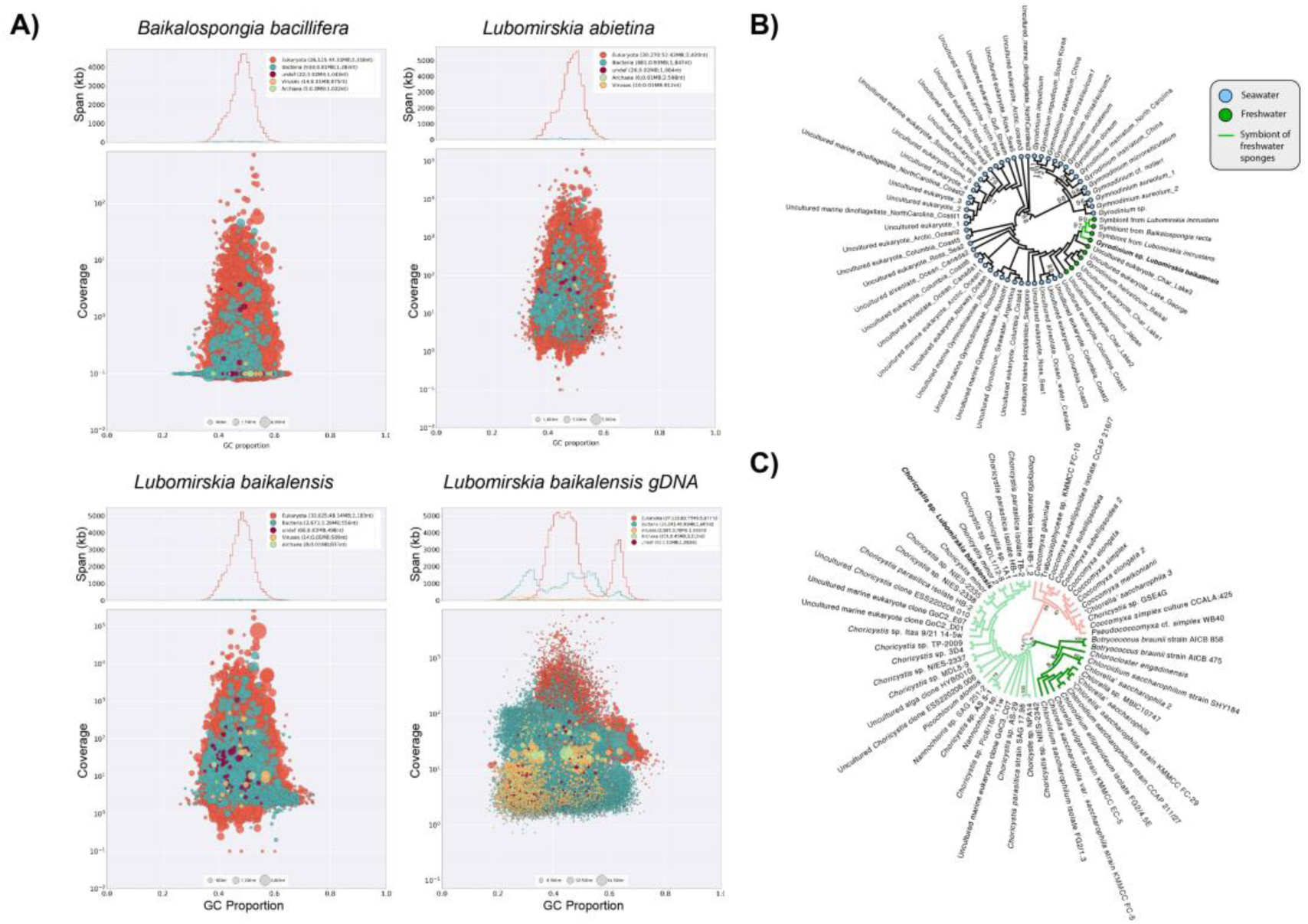
A) Blobplot results, showing distribution of annotated contigs according to GC content (x axis) and coverage (y axis) for the assemblies presented here. Summary statistics are also provided at the top of each panel. Note the bimodal distribution of GC content in the genome, which may represent differences between the coding and non-coding elements of the genome as observed in other species. B) Maximum likelihood tree showing relationship between *Gyrodinium sp* 18S rRNA sequence found in our *Lubomirskia baikalensis* sample and those identified previously. Note monophyletic group of freshwater samples, and internally to that clade, a monophyletic group of freshwater sponge symbiote sequences, as marked with symbols/colours noted in legend. C) Maximum likelihood phylogeny showing relationship of *Choricystis* sp. 28S-ITS1 sequence found in our *Lubomirskia baikalensis* sample compared to previously sequenced samples from Genbank. *Choricystis* samples shown at left (light green branches) with our sample shown in bold.

The quantity of this bacterial sequence data (or symbiont sequences) was, however, quite small, less than 1/50th of the quantity of sponge data for our transcriptomes, as can be seen most clearly in Supplementary File 3, which shows the distribution of our data by superkingdom and phylum. The proportion of bases mapped to Bacteria in our genomic sequencing was higher, around 5%, and may well consist of symbiont sequences of utility to future studies, and discussed further below.

Summary statistics relating to genome assemblies can be seen in Supplementary File 1, Table 3. A number of assembly algorithms were assessed for their utility, with SPAdes being the best performed by several metrics. Assemblies, including preliminary and alternate forms, have been uploaded to Figshare, with DOI: and URL: 10.6084/m9.figshare.6819812, https://figshare.com/s/fe36239c32bbf7342756.

We utilized GenomeScope to gain an understanding of the coverage of our dataset, potential levels of non-sponge DNA, and to estimate the genome size of *L. baikalensis.* The results of this analysis can be seen in Supplementary File 2. The inferred genome size of *L. baikalensis,* between 558 and 565 million base pairs in size, is considerably larger than that of many previously published sponge genomes (Supplementary File 1, Table 4). At the same time, the mitochondrial genome of this sponge is the largest found in demosponges (Lavrov et al 2012). The genome of *Amphimedon queenslandica* was estimated to be approximately 167 million base pairs in haploid size (Srivastava et al 2010) and *Tethya wilhelma* possesses approximately 125 million base pairs (Francis et al 2017). Estimates of 75 sponge species using traditional techniques (Jeffrey et al 2013) ranged from 0.04–0.63 pg in haploid size, or approximately 40-630 megabases. Smaller genomes were much more common, with only 6 genomes larger than 400 megabases. Our genome is therefore definitely larger than the average sponge genome. The freshwater sponges in that latter dataset, three members of the Spongillida, ranged in size from 0.31 to 0.36 pg (310-360 megabases). The size of *L. baikalensis’* genome is therefore not solely due to a freshwater environment, although this may play a part in its larger size. The unique demographic history and environment of Lake Baikal, and the unique evolutionary history of the lineage leading to *L. baikalensis,* will have played roles in the expansion of the genome of this sponge, in ways that will only be understood through examination of datasets such as the one presented here.

To estimate how much of the coding set of genes could be present in our genomic assembly, we used BLASTN megablast (-evalue 0.000001 -num_threads 8 -max_target_seqs 1-outfmt 6) to ascertain how many of the contigs present in our *L. baikalensis* transcriptome were present in the genomic assembly. 77,717 of the 81,951 contigs in the *L. baikalensis* transcriptome possessed a hit in the genome at this stringency. This strongly suggests that the transcribed cassette is well-represented by the genomic assembly.

### Annotation of assemblies

To test the content of both our genomic and transcriptomic assemblies, we used the BUSCO (“Basic Universal Single Copy Orthologue”) complements of highly conserved genes. In particular, we compared our datasets to the metazoan and eukaryote BUSCO complements, consisting of 978 and 303 genes respectively. The recovery of these gene families in our datasets was almost complete (Supplementary File 1, Table 5), and suggests that our transcriptomic assemblies are excellent resources for future work. Of the 303 genes in the eukaryotic BUSCO set, the maximum number of missing genes is 4 in our transcriptomic complements, and 17 in our genome assembly. Similarly, of the 978-strong metazoan BUSCO complement, the maximum number of missing genes is 46 in our transcriptomes, and 169 in our genome assembly. This compares favourably with the *A. queenslandica* genome. In the published cDNA set for *A. queenslandica*, 1.6% of the eukaryote set (5 genes) is missing, and 4.9% (49) of the metazoan complement. Our transcriptomic assemblies therefore are more complete in terms of gene recovery than that published resource, while our genome is almost as complete, despite its low read coverage depth and limited library complexity. Given this level of recovery of the BUSCO set, these assemblies are therefore likely to contain the vast majority of the genetic cassette of these species, with any genes expressed at the time of sampling likely to be recovered by our assembly.

We performed automatic annotation of the transcriptomic sequences detailed here using BLAST2GO, Interproscan and ANNEX. The complete annotations are attached to this manuscript as Supplementary File 4. In summary, of 80,829 total contigs, *Baikalospongia bacillifera* (A2) possessed 13,454 fully annotated contigs, while 58,890 contigs possessed no BLAST hit. For *Lubomirskia abietina* (A10), 93,299 contigs included 14,623 fully annotated sequences, with 69,862 had no BLAST similarity at the given thresholds. In *Lubomirskia baikalensis* (A8), these numbers were 81,877 contigs, 14,772 annotations, and 57,476 without significant similarity. The number of annotated contigs is therefore relatively stable between all of our samples, and as can be seen in Supplementary File 5, tend to be longer than those without significant similarity, as between 75 and 80% of the assembly (by nucleotide size) is incorporated into contigs with a BLAST hit.

It is important to note that novelties, such as novel genes specific to Lake Baikal sponges, will not be well described by this annotation process, which depends on similarity in the first instance for annotation to occur. For more information on novelty, please see the last section of this manuscript.

There is no clear signal of specific microbial sequence over-representation in our read data, which would be obvious at this point in the form of bacterial species with high numbers of “top hits”. However, some bacterial sequences are present, particularly that of the likely symbiont (Wilson et al 2014) *Candidatus* Entotheonella gemina, with 88, 76 and 85 hits respectively for our three transcriptomes. This species is therefore likely the most commonly present bacteria in our samples, or at least the one with sequences present in Genbank for assignment. Other bacterial species are also present, although less commonly. The fuller bacterial content of our assemblies is discussed in the next section of this manuscript.

Dinoflagellates are represented, albeit well down the list of commonly top-hit species, with *Symbiodinium microadriaticum* represented by 53, 66 and 51 sequences respectively. It is possible that some of the sequences without blast results represent dinoflagellate sequence, as the species noted to be symbionts of Lake Baikal sponges, such as those of the genus *Gyrodinium* (Annenkova et al 2011), are not well represented in the *nr* database.

Please note that we have not performed full gene prediction or annotation on the genome resource, as the relatively low contiguity and N50 of this resource (a consequence of low coverage and a lack of long read data) precludes most genes from occurring in their full length on a single contig. However, some information on the annotation of this resource can be derived from BLAST searches performed for Blobtools analysis, which incorporates an annotation step (see Fig 2 and Supplementary File 3). For full coding gene identification we recommend the use of our transcriptomes, while the genome will be of use for microsatellite identification, mitochondrial genomics, additional sequence determination and other research into the wider biology of these species.

### Bacterial Sequences

Our analyses revealed a small number of bacterial sequences were present in our datasets. Supplementary File 3 shows this clearly, with a maximum of 2.4% in our transcriptomes, and 5.44% in our genomic resource. The larger proportion of bacterial content in the genomic sample (SPAdes 500 bp+ assembly) is likely the result of poly A selection procedures in our transcriptomic samples, which will preferentially target eukaryotic mRNA.

We used the results of our Blobtools annotation pathway to understand the makeup of the bacterial sequence within our SPAdes 500 bp+ genome sample. The bacterial sequences that were present and identifiable in our *L. baikalensis* genomic sample had a relatively high abundance but low N50 when compared to eukaryote-annotated sequence, with more contigs (30,243 vs 27,115 annotated contigs) but a much smaller N50, 1,607 bp vs 5,877 bp. This indicates to us that bacterial sequences present represent a subsample of complete bacterial diversity. It is worth noting, however, that our assemblies are made with only a single sponge specimen, which may not be representative of the population as a whole.

Our *L. baikalensis* genome resource most commonly contained proteobacterial sequences (9,880 annotated contigs, correlating to 1.9% of the reads mapped to our assembly) but also contained more than 1000 contigs with similarity to Bacteroidetes, Verrucomicrobia, Actinobacteria and Cyanobacteria, in order of decreasing occurrence. Less commonly, Firmicutes, *Candidatus* Tectomicrobia (with a high N50, 3,788 bp), and Spirochaetes were observed in more than 100 contigs. All of these groups, with the exception of *Candidatus* Tectomicrobia, had a much lower N50 than the eukaryote (5,877 bp) and poriferan (5,088 bp) figures. This indicates that bacterial genomes were not inordinately represented in our data, and therefore were not well-assembled.

Our species top hit data revealed that *Candidatus* Entotheonella gemina was the most commonly hit individual species by BLASTP identity (phylum: *Candidatus* Tectomicrobia). Our Blobtools analysis, incorporating BLASTX, was able to categorise the number of contigs further. These results are available in full in Supplementary File 5. Previous studies of bacterial diversity in freshwater sponges (e.g. Gernert et al, 2005, Costa et al, 2013, Gaikwad et al 2016, Seo et al 2016) have found a range of bacterial sequences within members of the Spongillidae. Often there is overlap between species (e.g. Seo et al 2016) and in some cases freshwater sponge-specific lineages of bacteria can be found. For instance, freshwater-specific examples of Chlamydiae, Alphaproteobacteria, Actinobacteria and Bacteroidetes species were observed in Costa et al, 2013.

In *B. bacillifera* (A2) Proteobacteria were the most commonly observed bacterial phylum, with 392 contigs with N50 1,275. *Candidatus* Tectomicrobia was behind, with 157 contigs of 1,145 bp N50. In order from most common to least, Bacteroidetes, Firmicutes, Actinobacteria, Cyanobacteria and Verrucomicrobia were also present in at least double digit numbers of contig, the lattermost with an exceptionally high N50 (7,259 bp). Only 9 contigs of Chlamydiae were present, but their high N50 (9.999 bp) and the obligately intracellular nature of these make them worth noting. In *L. abietina* (A10) this result was mirrored, with 407 Proteobacterial contigs (N50 1,537 bp) and 203 from *Candidatus* Tectomicrobia (1,121 bp). However, in *L. abietina,* the order of commonality in other bacterial groups is (most -> least) Actinobacteria, Firmicutes, Bacteroidetes, Cyanobacteria, Verrucomicrobia (N50 7,270) and Chloroflexi. Only 8 Chlamydiae contigs were present, but the N50 of these, 10,037 was again striking. *Lubomirskia baikalensis* (A8) showed the same general trend, with one exception. Actinobacteria is the most commonly observed clade of bacteria in this sample, with 1,589 contigs of N50 297. Proteobacterial contigs (N50 1,087 bp) and *Candidatus* Tectomicrobia sp (191 contigs, 1,167 bp) are again well represented. Firmicutes, Bacteroidetes, Cyanobacteria, Verrucomicrobia (with a much smaller N50 than the other two samples, 976 bp) and Chlamydiae (N50 in this case only 1,618 bp) then round out the most common bacterial content in decreasing order of occurrence. Representatives of these phyla were identified previously in the *L. baikalensis* community (Kaluzhnaya et al 2012). It would be well worth using this data for targeted re-sequencing of further individuals of each of our target species to determine how representative these results are of bacterial symbiote diversity in the wider population.

This indicates to us that the content of our assemblies is consistent and coherent with the literature from other freshwater sponge species (e.g. Gernert et al, 2005, Costa et al, 2013, Gaikwad et al 2016, Seo et al 2016). The most likely individual bacterial symbiont species within our samples is clearly related to *Candidatus* Entotheonella gemina (Phylum: *Candidatus* Tectomicrobia). It is striking that this bacterial species made the transition to freshwater to join its host, although the contemporaneity of this has not been established here. The functions provided by this species therefore remain vital, even in the markedly differing freshwater environment (Wilson et al 2014, Liu et al 2016), and may be necessary for the survival of the host. This symbiote can protect against heavy metals, can supply energy in anaerobic environments and aid in CO_2_ fixation (Liu et al 2016), roles that would all be useful in freshwater in general, and Lake Baikal in particular. This is the first clear example of a sponge symbiont that is present in both marine and freshwater environments, a fact that clearly merits further investigation.

### Dinoflagellate, Symbiont and Other Sequence Content

*Symbiodinium microadriaticum* was represented by 53, 66 and 51 BLAST top-hits respectively in *B. bacillifera* (A2) *Lubomirskia abietina* (A10) and *Lubomirskia baikalensis* (A8). Of these hits, most were represented in two or more of the three species, with 7 represented in all three species. Only 3 (OLQ15661.1, OLP92806.1, OLP93934.1), 1 (OLP92903.1) and 2 (OLQ11623.1 OLP83851.1) sequences from the *nr* database were present in only one of the three species. The degree of overlap between the hits in our samples leads us to conclude that all species of sponge make use of dinoflagellate symbionts, as shown in Annenkova et al (2011), and that the same genes are expressed at high levels in all three species. All sequences and their alignments with searched sequence are available for download in Supplementary File 5.

It is likely that these sequences do not actually represent *S. microadriaticum* – instead, they match to that species as it is the best-sequenced symbiotic dinoflagellate within the *nr* database. This is borne out by the BLAST alignments present in Supplementary File 5. In no case are hits identical to known *S. microadriaticum* sequence. In the hit with the lowest *E* value, (to 132 kDa protein, OLP79640.1, *E* value 7.7e-130) there are 81 mis-matches over a 1058 amino acid alignment. These sequences are therefore likely from a related species of dinoflagellate, but not *S. microadriaticum* itself.

To determine which species these sequences could have been derived from, we checked for common molecular markers from dinoflagellates and compared these to genes of known orthology. We found a 18S sequence with 99.67% similarity to *Gyrodinium helveticum* in our *L. baicalensis* genomic sample, as shown in a phylogenomic context in Fig 2B. Our *Gyrodinium* 18S sequence is therefore further confirmed as a symbiont of Lake Baikal sponges (Annenkova et al 2011), and our phylogeny shows a monophyletic group of freshwater *Gyrodinium* species, that are the sister clade to free-living *G. helveticum*, suggesting that they may have entered that environment contemporaneously with their hosts. They almost certainly play a role in supplying their sponge host with nutrition through photosynthesis, and may supply specific metabolites to their host (Müller et al 2009). All accession numbers and alignments used in making this phylogeny can be found in Supplementary File 5.

We were also able to use our Blobplot results to examine the contents of our genomic resource. A total of 104 contigs, with an N50 length of 2,651 bp, were identified as belonging to the Bacillariophyta by blast similarity. This is not a high number, and indicates that the symbionts are not present in high numbers within our samples, at least at the time of sampling.

Many of the best assembled sequences present in our genomic data derive from unicellular algae, which are common symbionts of freshwater sponges (Feranchuk et al. 2018). In our genomic sample, the Chlorophyta are the second most commonly-hit phylum after Porifera, with 1,359 very well assembled contigs of N50 21,412 bp - an N50 fourfold higher than that for our sponge data. In our transcriptomes, 189 *L. baikalensis* (A8) but only 14 and 21 contigs in *L. abietina* (A10) and *B. bacillifera* (A2) respectively. As our genomic and transcriptomic sample for *L. baikalensis* are derived from the same tissue, we suspect that a member of the Chlorophyta was particularly abundant in that specimen, and could have been performing a symbiotic role, as has been reported previously in freshwater sponges (Sand-Jensen & Pedersen 1994; Feranchuk et al 2018). They play clear roles in providing sustenance through photosynthesis, which greatly benefits the sponge host (Chernogor et al 2013). Phylogenetic analysis of the 18S-ITS1 sequence of this symbiont from our genomic sample (Fig 2C) identifies it as *Choricystis* sp., placed as sister to a free-living *Choricystis minor* collected from a lake in Germany (Krienitz et al. 1996). These algae are known symbionts of freshwater sponges, and this record confirms their role in Lake Baikal sponges. The accession numbers and alignments of sequences used for this phylogeny are available in Supplementary File 5.

We also note, as mentioned in the bacterial section above, bacterial species with similarity to *Entotheonella* (Wilson et al 2014) are also present. These findings, alongside those published previously, suggest the microbial symbiota of the sponges of Lake Baikal is diverse and species dependent. As noted above, re-sequencing multiple individuals from a range of seasons, using the data presented here to aid primer design, would allow more complete understanding of the full contribution of symbionts to the survival of these species.

### Mitochondrial Genomes, and Mitochondrial and Nuclear-derived Phylogenies

The mitochondrial genomes of several Lake Baikal sponges have been published previously (Lavrov 2010, Lavrov et al 2012, Maikova et al 2016). They display a number of special characteristics, notably the proliferation of small “hairpin” inverted repeats, which act as a source of variation. To confirm the identity of the sponges sequenced here, and to identify regions of difference at the DNA and coding level, we studied the sequence of the mitochondrial genomes of the three species investigated, and compared them to previously published work. Of the three species included in this manuscript, only *L. abietina* has not been sequenced previously. *L. abietina* sequences have been uploaded to Genbank with accession numbers MH697685-MH697697.

Of the sequences presented here, the mitochondrial sequence of *L. baikalensis* derived from sample A8 seems closely related to that of the previously sequenced example of this species, while *B. bacillifera* differs somewhat from the example in Genbank, which appears more similar to *B. intermedia.* The small number of changes between these sequences, however, may render the differences inconsequential compared to morphological evidence (as suggested by low posterior probability of a real clade occurring between these sequences, which could be collapsed to a polytomy), and indels also add data, as discussed below. The phylogeny shown in Figure 3 seems to indicate paraphyly in both *Lubomirskia* and *Baikalospongia*, albeit in some cases with limited posterior probability support. This mirrors the phylogeny previously observed in Lavrov et al 2012 (Fig 6 of that paper), adding extra taxa to the outline presented there. *Swartschewskia papyracea* and *Rezinkovia echinata* seem to render the *Lubomirskia* paraphyletic, while the *Baikalospongia* seem to be paraphyletic as a mixed paraphyletic sister group of this clade. From the evidence presented in this figure, taxonomic revision of these clades appears warranted.

**Figure 3:**
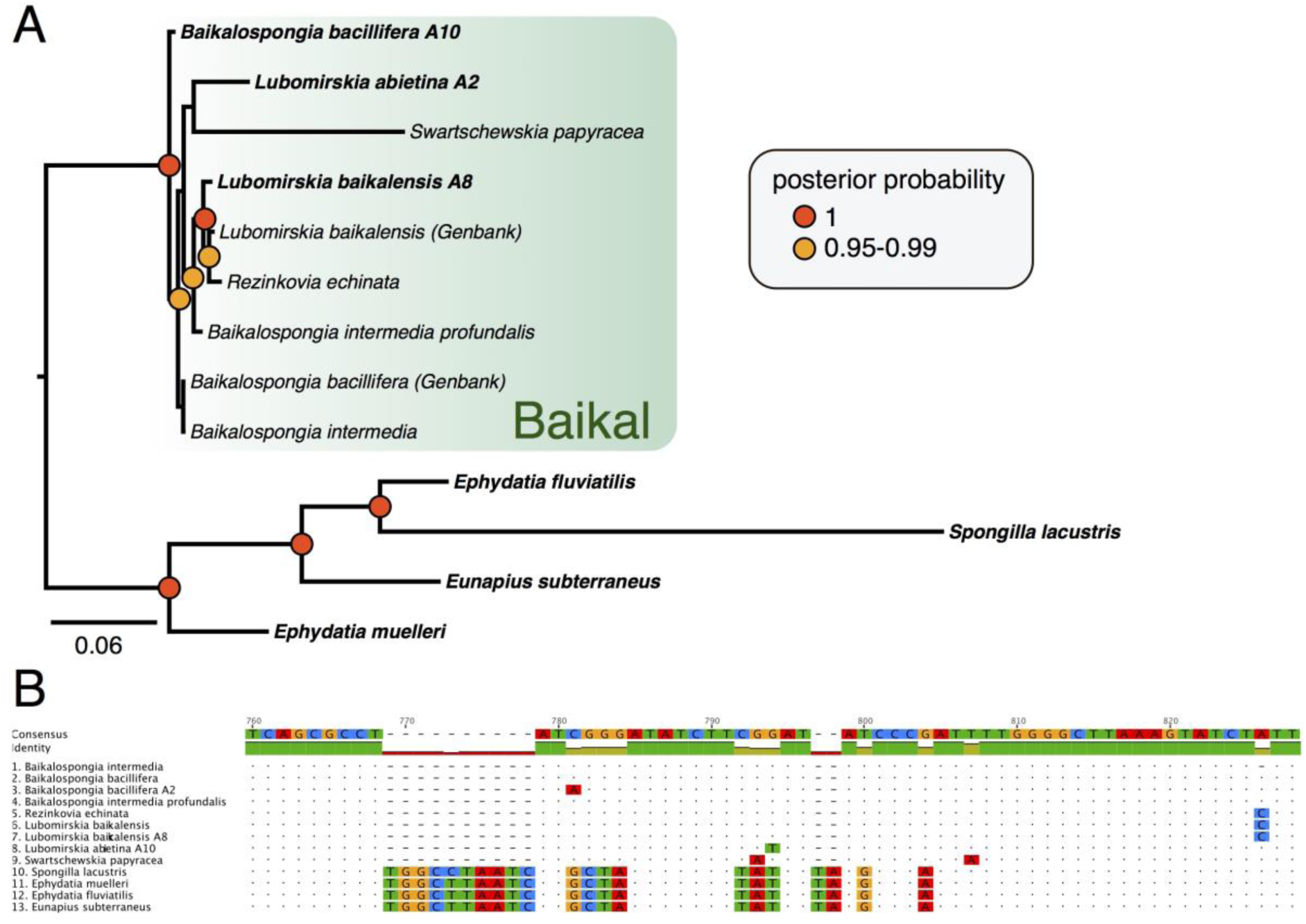
A) Phylogenetic relationships of a variety of freshwater sponge species, inferred using Bayesian methods, based on alignment of nucleotide sequence from mitochondrial protein coding and rRNA genes. Numbers at bases of nodes indicate posterior probability, number at base of tree (under scale bar) indicates number of changes per site at unit length. B) example of indels not used in tree, but nonetheless present in alignment. These indels (example from *Nad1* provided) provide further data that can be used to support clades, but are not represented in the phylogeny shown (as any site without sequence data is excluded from that analysis). Sponges endemic to Lake Baikal are, on the basis of this evidence, firmly supported as a monophyletic clade, to the exclusion of other freshwater sponges.

**Figure 6:**
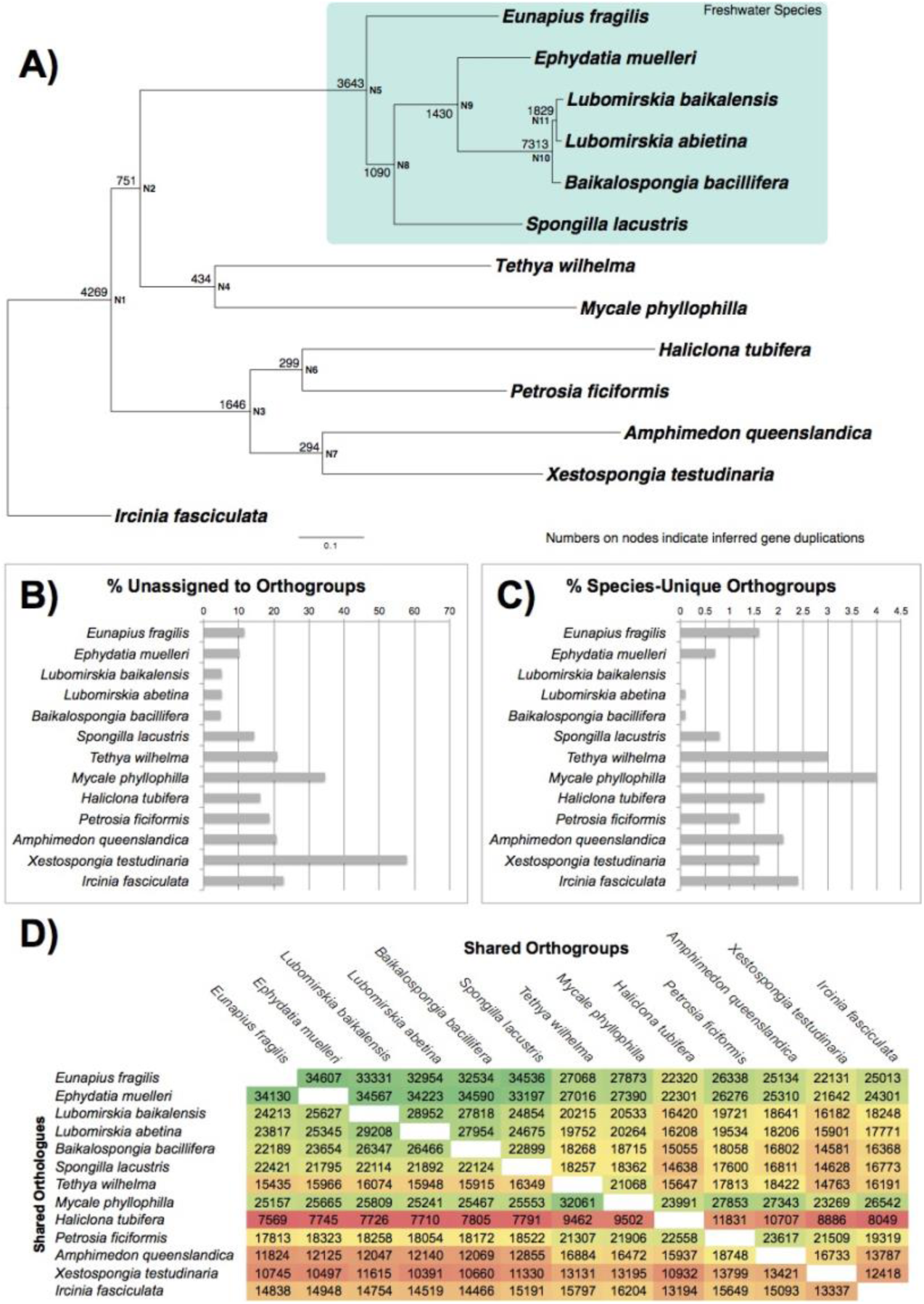
Analysis of novelties present in freshwater sponge transcriptomes when compared to a range of other genomes and transcriptomes. A) Phylogeny of freshwater sponges together with marine outgroups used in this analysis. Tree is rooted with *Ircinia fasciculata.* Mapped onto the phylogeny at bases of nodes are the number of duplications that were inferred to map to each node. B) Contigs unassigned to orthogroups (and thus unique, and single copy, within individual species). C) orthogroups present only in a single species (and thus unique to the species, but with duplication/alternative isoforms present). D) Matrix of numbers of orthogroups (top) and orthologues (bottom) shared between species, coloured as heat map, with green showing the highest number and red the lowest numbers for each comparison. These numbers are given for for every species pair, and can be read by finding the names of the species for comparison on the x and y labels, and moving to the site of overlap.

The phylogeny shown in Figure 3, and particularly the branch lengths, does not fully show the level of divergence of the Lake Baikal clade, as gaps and indel regions have been excluded for the purpose of phylogenetic analysis. Several of these sites show clear diagnostic differences between the sponges of Lake Baikal and other freshwater species. One of these regions, that of *nad1*, can be seen in Figure 3B, by way of example. This alignment clearly shows the disparity between freshwater sponges of Lake Baikal and those from further afield, but also (in position 826) shows how a diagnostic difference separates *L. baikalensis* and *R. echinata* from the other species examined here. A fuller examination of the poriferan diversity of Lake Baikal, considering both morphological and genetic evidence, is therefore likely warranted to resolve these species into a taxonomically and systematically cohesive framework.

As part of the process of studying the signatures of selection within freshwater sponges, the Phylotreepruner output alignments of all 3,222 genes to be tested were concatenated, and used to construct a robust phylogeny for our species, spanning 3,242,264 amino acid residues although all gaps and indels were removed for phylogenetic inference. This phylogeny (and our tests of signatures of selection) included the three species of Lake Baikal examined here, alongside three other members of the Spongillida (*Spongilla lacustris, Eunapius fragilis* and *Ephydatia muelleri*). Five marine outgroup species (*Mycale phyllophilla. Haliclona tubifera, Petrosia ficiformis, Amphimedon queenslandica* and *Ircinia fasciculata*) were chosen for their taxonomic position and sequenced gene complements.

Phylogenetic inference was performed by MrBayes using a by-gene partitioned GTR analysis, and the resulting phylogeny can be seen in Figure 4A. In this phylogeny, the freshwater sponges form a distinct clade corresponding to the order Spongillida (Manconi and Pronzato, 2002), with maximal posterior probability support. *Mycale phyllophila* is noted as the outgroup to this clade, suggesting poecilosclerids are more closely related to Spongillida than the other species represented here, albeit with weak (0.72) posterior probability. The three species of haplosclerid examined form a monophyletic group with posterior probability of 1, and the sole member of the Dictyoceratida (and Keratosa) present in our sample, *Ircinia fasciculata*, is used to root the tree shown here. This is in agreement with the view of inter-relationships put forward in Morrow and Cárdenas 2015 (see Fig 2 of that work).

**Figure 4:**
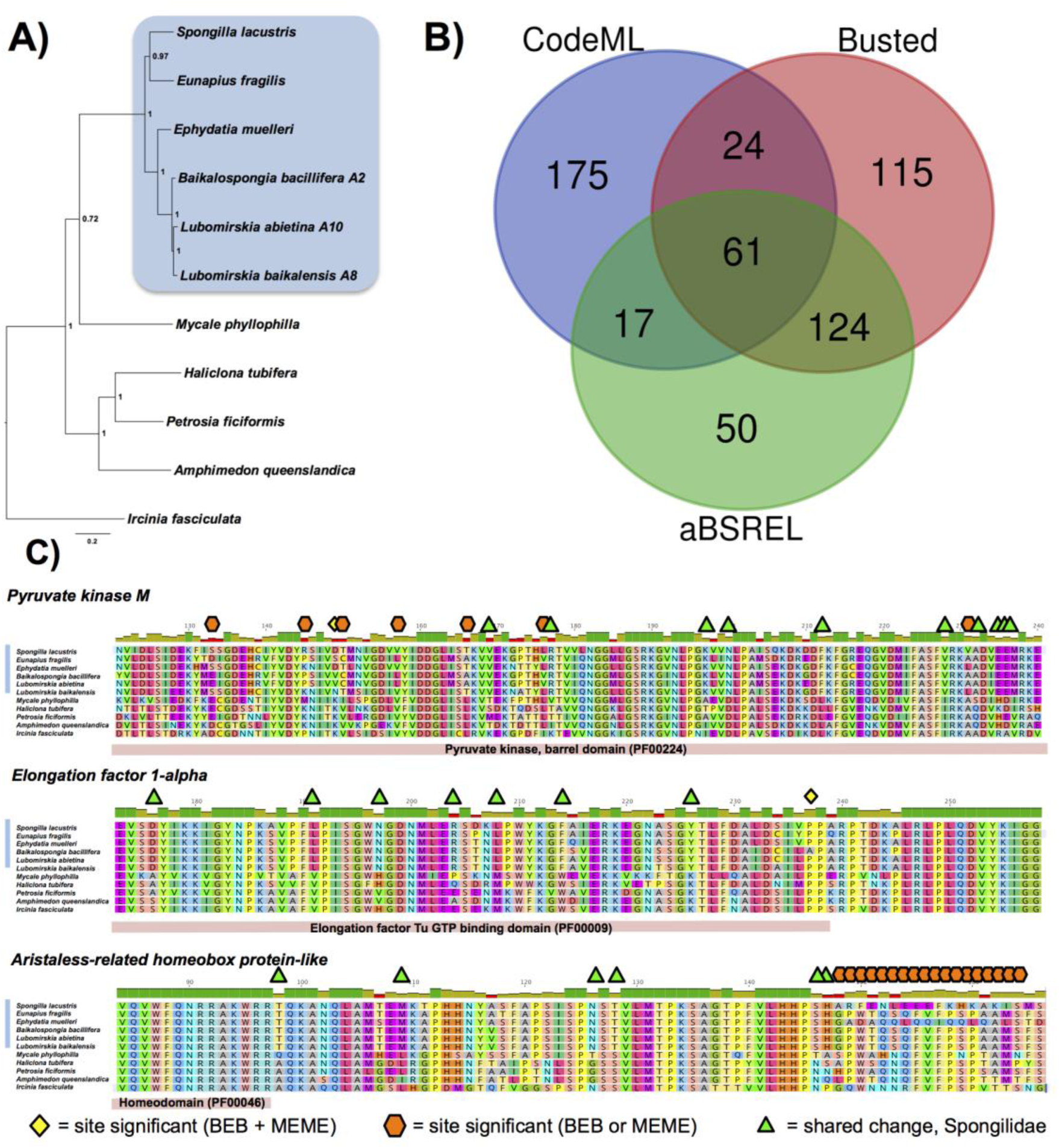
A) Phylogenetic relationship of freshwater sponge species and outgroup taxa, inferred from a concatenated multiple gene alignment using partitioned GTR analysis in MrBayes (3,222 genes, 3,242,264 site alignment, before trimming). This tree was used as the basis for tests of selection on the freshwater lineage, the results of which are summarised in: B) Orthogroups displaying significant results for tests of selection under several tests, shown in a Venn diagram indicating consilience of results. C) examples of genes and sites under selection, with sites marked according to key at bottom of figure. Domain locations are indicated below each gene as appropriate. Note that further site-level (MEME, BEB) results are available in Supplementary File 5.

Within the Spongillida, *Lubomirskia* forms a monophyletic grouping, with *Baikalospongia* as the sister clade. This is in contrast to the mitochondrial tree, and likely represents the increased depth of data available to distinguish these clades. *E. muelleri* is strongly supported as the outgroup to the Lubomirskidae, with *E. fragilis* and *S. lacustris* placed in a monophyletic group as sister to the (Lubomirskidae + *E. muelleri*) clade. This mirrors the results of prior, individual and several gene/ITS based phylogenetic studies in freshwater sponges, e.g. (Meixner et al 2007, Erpenbeck et al 2011). As such, the monophyly of freshwater sponges, and the hypothesis of the independent evolution of endemic sponge species locally, from cosmopolitan founder species, seems secure (Meixner et al 2007).

### Signatures of Molecular Adaptation to Freshwater Conditions

To discern how freshwater sponge species have evolved to cope with their environments, we have utilized the Hyphy and PAML software suites to identify genes where there are marked signals of selection in freshwater sponges when compared to the ancestral condition. The phylogeny of sponges used to conduct these tests, the concatenated nuclear gene phylogeny, can be seen in Figure 4A, and is discussed further above. In total, 3,222 single copy orthologous alignments remaining after pruning paralogous sequences with Phylotreepruner, henceforth referred to as “orthogroups”, were tested from the 11 transcriptomes we used in this experiment.

We used the concilience of several tests to validate our findings, to avoid artifactual or spurious results. In particular, we used CodeML, Busted and aBSREL to test for signatures of selection using a combination of approaches (branch-site model, alignment-wide episodic diversifying selection, and adaptive branch-site random effects likelihood, respectively). These methods use slightly varying means to detect selection, so a combinatorial approach is most suitable for ensuring that identified selective events are well supported by evidence, rather than the product of branch-specific signals or errors that could yield false positives (Venkat et al 2018).

A total of 277, 324 and 252 orthogroups were identified as bearing signatures of selection by CodeML, Busted and aBSREL respectively. These represent 566 unique orthogroups, with 287 of them overlapping in some way, as can be seen in Figure 4B. The fact that 566 orthogroups, 17.6% of those tested, show some sign of selection is suggestive of broad scale changes in sequence as a consequence of adaptation to a freshwater lifestyle. Of these, 61 orthogroups were identified by all 3 of our tests (see Figure 4B), and it was these that we concentrated on in further analyses. These orthogroups, which can be seen listed in Table 1, with full details given in Supplementary File 5, represent a variety of gene families. Of these 61 sequences, only one was un-annotatable by BLAST identity to previously described genes. Annotation was aided by the inclusion of the most recent *A. queenslandica* resource in our test dataset, and the sole unannotatable orthogroup represents a newly identified sequence (in the recent *A.queenslandica* transcriptome) not originally annotated in the *A. queenslandica* genome, which nonetheless is present in all of the sponge species studied in this test. It does not have any matches on the NR or PFAM-A databases, with an *E* value cutoff of 1, It contains an ORF 175 amino acids in length, and is thus likely to be a *bona fide* protein coding gene.

**Table 1:** Orthogroups (OG) with consilient signals of molecular evolution. 35 of these could be mapped to Gene Orthology (GO) Terms, as described in text. Full details of sites and significance, along with mapped GO terms can be found in Supplementary File 5.

| Alignment Number | Genbank Hit | Mapped to GO Term(s) | Alignment Number | Genbank Hit | Mapped to GO Term(s) |
| --- | --- | --- | --- | --- | --- |
| OG0000044 | fibrinogen C domain-containing protein 1-A-like | No | OG0001408 | uncharacterized protein LOC109593314 | Yes |
| OG0000120 | zinc transporter 2-like | Yes | OG0001488 | UPF0600 protein C5orf51 homolog | No |
| OG0000141 | neurobeachin-like | No | OG0001657 | protein ADP-ribosylarginine hydrolase-like | No |
| OG0000142 | actin-3-like | No | OG0001662 | elongation factor 1-alpha | Yes |
| OG0000150 | proteinase T-like | Yes | OG0001704 | apoptosis-inducing factor 3-like | No |
| OG0000161 | autophagy-related protein 16-1-like | No | OG0001826 | WD and tetratricopeptide repeats protein 1-like isoform X1 | No |
| OG0000227 | uncharacterized protein LOC109592610 isoform X2 | No | OG0002175 | tyrosine-protein phosphatase non-receptor type 11 isoform X2 | No |
| OG0000236 | protein F37C4.5-like | No | OG0002186 | U3 small nucleolar ribonucleoprotein protein MPP10-like | Yes |
| OG0000300 | dual oxidase 1-like | Yes | OG0002195 | rho GTPase-activating protein 26-like isoform X2 | Yes |
| OG0000375 | SEC14-like protein 2 | Yes | OG0002335 | SH3 domain-containing kinase-binding protein 1-like isoform X2 | Yes |
| OG0000378 | glutamyl aminopeptidase-like isoform X2 | Yes | OG0002484 | radixin-like | Yes |
| OG0000422 | annexin-B12-like, partial | No | OG0002622 | tetratricopeptide repeat protein 14 | No |
| OG0000434 | long-chain-fatty-acid--CoA ligase ACSBG2-like | No | OG0002646 | alpha-xylosidase-like | Yes |
| OG0000519 | gamma-glutamyltranspeptidase 1-like | Yes | OG0002712 | uncharacterized protein LOC109586818 | No |
| OG0000528 | dnaJ homolog subfamily A member 3, mitochondrial-like isoform X1 | Yes | OG0002718 | pyruvate kinase PKM-like | Yes |
| OG0000631 | cytochrome b-245 heavy chain-like | Yes | OG0002786 | heparan-alpha-glucosaminide N-acetyltransferase-like isoform X2 | No |
| OG0000658 | disintegrin and metalloproteinase domain-containing protein 17 | Yes | OG0003043 | uncharacterized protein LOC100635535 | No |
| OG0000696 | 2'-5'-oligoadenylate synthase-like protein 2 | Yes | OG0003137 | uncharacterized protein LOC105316736 isoform X2 | Yes |
| OG0000757 | protein vav-1-like | Yes | OG0003269 | Acyl Co-a binding | No |
| OG0000774 | TGF-beta-activated kinase 1 and MAP3K7-binding protein 1-like | Yes | OG0003737 | tetratricopeptide repeat protein 1-like isoform X1 | No |
| OG0000885 | tyrosine-protein kinase Tec-like | Yes | OG0003883 | flowering time control protein FY-like | No |
| OG0000894 | poly(rC)-binding protein 2-like | No | OG0004236 | dual oxidase maturation factor 1-like | Yes |
| OG0000984 | ataxin-7-like protein 3 | No | OG0004507 | stromal membrane-associated protein 1-like | Yes |
| OG0001031 | integrin alpha-9-like | Yes | OG0004509 | brefeldin A-inhibited guanine nucleotide-exchange protein 1 | Yes |
| OG0001052 | glucosidase 2 subunit beta-like, partial | Yes | OG0005056 | homocysteine-responsive endoplasmic reticulum-resident ubiquitin-like | Yes |
| OG0001078 | protein DD3-3-like | No | OG0005<br>907 | protein tweety homolog 2-like | Yes |
| OG0001082 | guanine nucleotide-binding protein G(q)<br>subunit alpha-like | Yes | OG0005<br>914 | UNC93-like protein MFSD11 | Yes |
| OG0001093 | probable ATP-dependent RNA helicase<br>DDX49, partial | Yes | OG0006<br>570 | aristaless-related homeobox protein-like | Yes |
| OG0001117 | rho GTPase-activating protein 39-like<br>isoform X1 | Yes | OG0007<br>030 | 28 kDa heat- and acid-stable phosphoprotein-<br>like | No |
| OG0001194 | importin subunit alpha-6-like | Yes | OG0007<br>208 | pre-mRNA-processing factor 40 homolog B-like | No |
| OG0001217 | No BLAST/PFAM Annotation | No |  |  |  |

The 61 orthogroups highlighted by this analysis correspond to transcription factors, structural proteins, membrane transport molecules and a variety of other gene families (Table 1). This diversity reflects the wholesale nature of changes required by adaptation to a novel freshwater environment. To highlight some genes in particular, transporter genes such as *zinc transporter 2* (OG0000120) and *importin subunit alpha-6* (OG0001194), transcription factors such as *aristaless-related homeobox* (OG0006570), structural peptides such as *actin 3* (OG0000142) are all implicated. However, despite this diversity, some genes have clear roles in homeostasis and membrane transport that would be under particular pressure as a result of the adaptation to a freshwater environment. Several of the genes identified by our analysis are transmembrane or membrane-associated proteins, such as the aforementioned transporters, *integrin alpha-9* (OG0001031), *neurobeachin* (OG0000141), *UNC93-like protein* (OG0005914) and *tweety homolog 2 (*OG0005907). Several are also involved in the Rho GTPase pathway, including *vav-1* (OG0000757), *rho GTPase-activating protein 39* (OG0001117) and *rho GTPase-activating protein 2* (OG0002195). Given the widespread roles of Rho GTPases in maintaining homeostasis and cellular functionality (Bustelo et al 2007) their modification to a freshwater environment under selective pressure would be necessary.

We investigated the nature of the precise changes, and the exact sites under selection pressure, in these sequences by performing tests in MEME (Mixed Effects Model of Evolution, Murrell et al 2012, 2015), as well as by investigating the BEB (Bayes Empirical Bayes, Yang et al 2005) results from CodeML analysis. Of the 61 orthogroups identified as significant by all 3 tests described above, all but one possessed at least one amino acid site noted as under significant evolutionary pressure by either our BEB or MEME test. The sole exception, OG0002186, is annotated to be *U3 small nucleolar ribonucleoprotein protein MPP10-like*. These results are also listed in Supplementary File 5, although please note that site number refers to that position in the alignment, and not an individual amino acid (as this will vary from species to species). This extra, almost perfect, support from site-based tests gives us even more confidence in the veracity of these results, and the strength of the selection pressures brought to bear on these genes.

In Figure 4C we show some examples of the sites that are flagged by MEME and BEB as under selection. Three disparate genes are shown, *Pyruvate kinase M, Elongation factor 1-alpha*, and *Aristaless-related homeobox protein*. These were chosen for their interesting patterns of selection under BEB and MEME analysis, as well as their diverse functional utility.

*Pyruvate kinase M* catalyzes glycolysis, allowing energy production even in anoxic environments (Gupta and Bamezai 2010). In this gene there are a raft of changes identified by either BEB or MEME as significant proof of positive selection, but in only one case (site 149) do these tests agree. However, these sites do cluster in close proximity. In *Elongation factor 1-alpha* (Sasikumar et al 2012), which aids in delivery of tRNA to the ribosome, there are numerous freshwater sponge-specific changes, which are shared across the monophyletic Spongilidae. However, these are not necessarily detected as significant changes by BEB and MEME analysis. Instead, a change from proline (with its large cyclic sidechain) to alanine (with its small aliphatic sidechain) in *B. bacillifera* is noted by both tests as the most significant change to have occurred in the sequence of this gene.

The last example, *Aristaless-related homeobox protein*, a member of the PRD class of homeobox-containing genes, has a cluster of sites identified as positively selected, lying together towards the C terminus of the protein from the homeodomain. These sites are detected by MEME and BEB, but again, instead of agreeing on the same sites, only one of these two tests flags positive selection at any individual site, in an interleaving fashion. The exception to this is only at site 173, where they agree. It is clear from the alignment shown in Figure 4C that this region of the protein is more prone to change than the remainder of the protein sequence. This conservation is especially clear in the indicated homeodomain region. The potential for positive selection in the variable region is clear, although the role of this section of the protein is at present unknown.

We examined our BEB results to see whether some amino acids were more likely to be the site of selection than others. Of the 424 sites identified by BEB across the 61 orthogroups, the most likely amino acid to exhibit positive selection is lysine (K), which occurs 38 times. Serine (S) occurred 33, leucine (L) 29, and aspartic acid (D) and alanine (A) occurred 28 times. These amino acids contain a variety of charged, polar and hydrophobic side chains, and thus these changes are likely to reflect a variety of changes within these molecules. In contrast, tryptophan (W) was the least likely to be positively selected, occurring only 4 times, likely a result of the large side chain which would prove difficult to incorporate into protein structures. Cysteine (C), 7, and phenylalanine (F) and tyrosine (Y), with only 10 occurrences, were similarly less likely to be subject to positive selection, likely as a result of their tendency to form cross links and large side chains respectively.

The large number of changes and the diversity of the genes affected reflect the profound changes which must have occurred across the genome of freshwater sponges in order to adapt to the markedly different osmotic, climatic and environmental differences posed by freshwater environments. As all freshwater sponge species examined here are descended from the same common ancestor, we are unable to use convergent signal to verify that these signals of selection are the result of freshwater specific cues. They could therefore derive from other selection pressures within this clade in the time since their stem lineage diverged from their sister taxa. However, as these freshwater sponges share these signatures of positive selection, they are likely to be necessary for their survival to the present day. Functional testing of these proteins and their performance in freshwater when compared to their ancestral forms would allow the precise nature of the conformational and functional changes in these proteins to be determined, but is beyond the scope of this manuscript,

Previous “omic” studies looking into adaptation to freshwater environments have tended to focus on metazoans with complex body plans, for example fish (Mäkinen et al 2008) and prawns (Rahi et al 2017). The molecular adaptations of these species to less saline environments have often been found to be focused on genes expressed in gills and kidneys, as these are the organs most involved in maintaining homeostasis under the contrasting osmotic pressure found there. Sponges must utilise such adaptations across the entirety of their bodies, without the benefit of impermeable membranes, shells or skins. It is unsurprising that sponges exhibit profound changes across the majority of their molecular repertoires as a result of the pressures of freshwater environments. As sponges only populated freshwater on a single occasion, despite being widespread in marine environments, this transition cannot have been straightforward. Studies such as those described here will allow us to understand the myriad demands of this process, and contrast them with those encountered by independent transition events to the same environment.

### Over-representation of Gene Families or Categories

To gain deeper understanding of the links between the seemingly disparate genes with evidence for selection, contrasted with those genes without such evidence in freshwater conditions, we used the established system of gene ontology (GO) categories. This allowed us to investigate the links between our gene lists by determining how they are linked by shared molecular functions and pathways. GO categories over-represented in the 61 positively-selected genes seen in Table 1 were annotated for all three sub-ontologies (biological process, cellular component, molecular function) within the GO schema, and the significantly over-represented GO terms can be seen in Fig 5A. 35 of the 61 genes were able to be mapped to GO terms (Table 1), and often these were annotated in more than one sub-ontology. It should be noted that not all genes are included in the GO database, and absence of GO term does not mean that a gene is a novel or uncharacterised.

**Figure 5:**
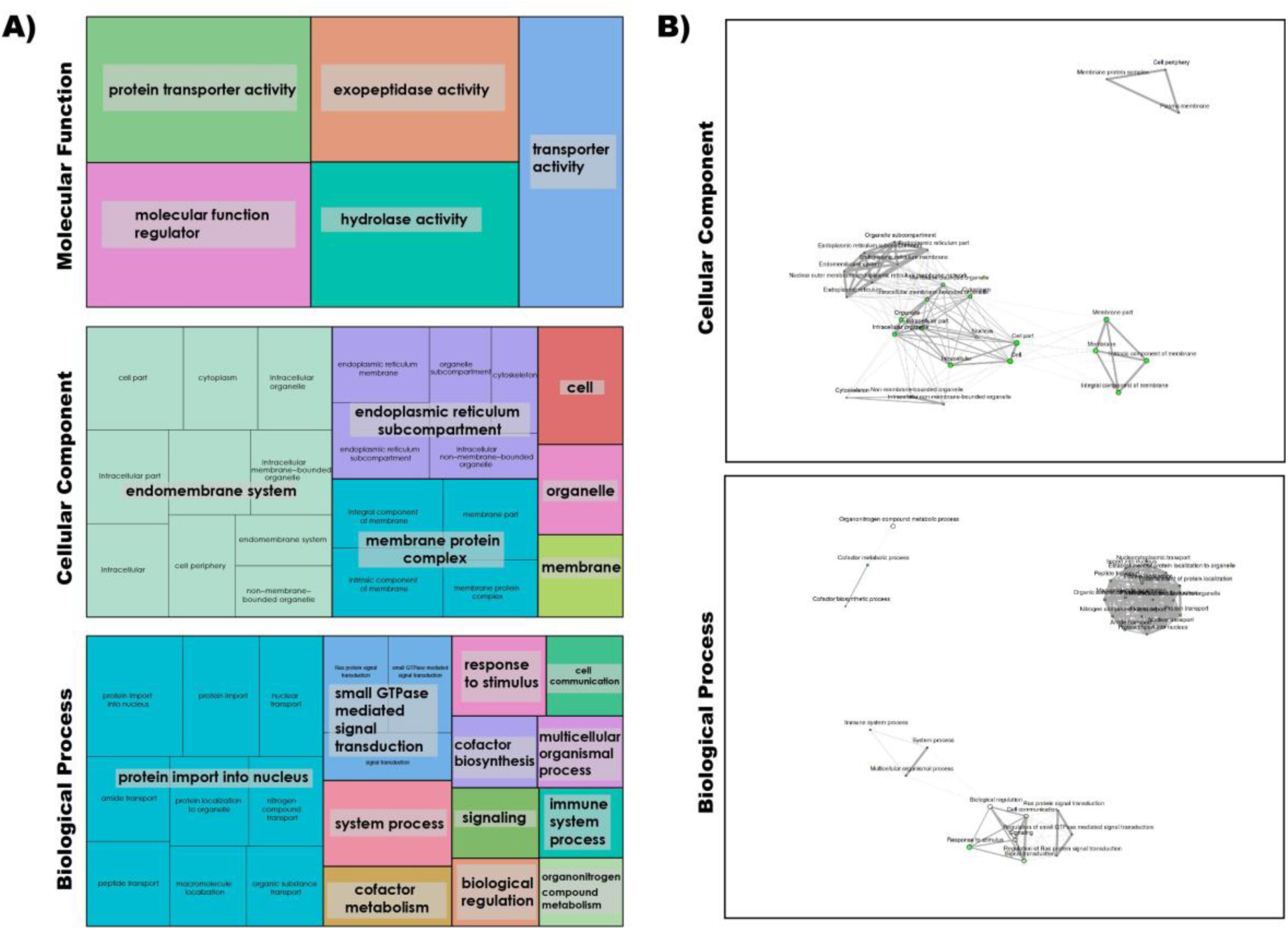
A) GO categories enriched (P-value cutoff (FDR) 0.05) in the 61 genes identified as under positive selection by all tests performed, as displayed by REVIGO. These are displayed according to GO sub-category (Molecular Function (19 annotated genes), Cellular Component (20 genes) and Biological Process (23 genes)), with area of coloured squares corresponding to the proportion of GO terms represented. B) ShinyGO network analysis of enriched GO terms for Cellular Component and Biological Process sub-categories (P-value cutoff (FDR) 0.05) within the 61 genes with consilient evidence for positive selection listed in Table 1. Nodes each represent an enriched GO term, with size corresponding to number of genes with this term. Lines connect related GO terms, and line thickness reflects percentage of genes that overlap for these categories. Please note in particular degree of overlap and connectedness of these categories.

Fewer significantly over-represented categories were seen in the Molecular Function (MF) sub-ontology (Fig 5A) than in the other two sub-ontologies. Both transporter activity in general and protein transporter activity in particular are noted as over-represented, and likely reflect changes in how these molecules function in freshwater conditions. Changes in transporters are often observed in transitions between marine and freshwater environments (e.g. Alverson 2007, DeFaveri et al 2011). ‘Molecular function regulation’, exopeptidase activity and hydrolases are all similarly over-represented in genes under selection, suggesting that life in hypotonic solutions poses problems to these functions within the cell, and requires widespread changes.

The Biological Process (BP) and Cellular Component (CC) categories contain more diversity than the MF category, and in particular many genes are mapped to specific roles within these GO ontologies (smaller text within figures, Fig. 5A). These can be related directly back to the identity of genes listed in Table 1 (with all GO mapping provided in Supplementary File 5). Genes such as *zinc transporter 2-like* have clear affinity to membranes and homeostasis, and are well mapped to GO terms matching such roles in freshwater species.

It is clear to see how these GO categories share commonalities between BP, CC and MF sub-ontologies. Membranes feature prominently in our CC figure (with membrane, endomembrane system, membrane-protein complex, and endoplasmic reticulum subcompartment GO terms listed). Transport (and particularly protein import), signalling and communication GO terms are prominent in our BP figure. Together with previously discussed MF results, membranes, transport and communication are key links between the GO terms of the genes under selection.

Figure 5B shows these enriched GO terms represented as networks for the CC and BP sub-ontologies (MF categories did not overlap meaningfully, and are not shown). These networks show that the enriched GO terms within our genes, for all their apparent disparity when viewed as a list, overlap considerably in componentry and biological process. Within total ‘GO space’, enriched GO terms are tightly clustered for both CC (upper) and BP (lower) networks. Links can be seen between the disparate GO terms listed in Fig 5A, with only 2 (CC) and 3 (BP) separate clusters in our dataset.

GO analysis therefore suggests that within our gene list (Table 1) there is coherent, linked GO signal, and the list, while containing genes seemingly unrelated at first sight, is linked by underlying processes. Freshwater sponges will, at the cellular level, be under similar evolutionary pressures to other organisms who have made the move to this environment. It is therefore both expected and confirmation of the validity of our analyses that genes and mechanisms involved in membrane function, transport and related processes, which would be intimately affected by changes in osmotic pressure and environment, appear in our GO terms and networks.

### Novelty in Freshwater Sponge Gene Complements

We assayed for novel genes and gene family expansions to discern whether there was any systematic change in the occurrence of novelties in freshwater sponges. The results of these investigations can be seen in Figure 6, as inferred by Orthofinder 2, which infers orthogroups using a combined reciprocal search (DIAMOND), clustering and phylogenetic approach, through generation and interpretation of gene trees. These orthogroups do not always correspond to individual genes, due to effects such as paralogy, but consist of “all sequences descending from a single gene in the last common ancestor of all the species being considered” (Emms and Kelly 2015).

We mapped the occurrence of duplications within orthogroups within our dataset, and displayed them on a species tree (made from our data, concatenated and partitioned) in Figure 6A. As can be seen from this data, duplication is particularly common at node N10, leading to Lake Baikal sponges. 7,313 duplications are noted here, and a further 1,829 in the node leading to the genus *Lubomirskia.* Generally, high levels of duplication are seen throughout the freshwater sponge clade when compared to marine outgroups. This is not due to the use of transcriptomes rather than genomic sequence, as transcriptome data are also used for 4 of the 7 marine species and do not show this trend. Only at N1, the base of the Heteroscleromorpha (*sensu* Cárdenas et al 2012) are similarly high levels of duplication found (4,269 duplications). A moderate signal for duplication is also seen at N3, at the base of the Haplosclerida (1,646 duplications), but this not comparable to the large degree of duplication seen in the Spongillidae (for example, 3,642 duplications at N5, at the base of the Spongillidae). Of the 61 genes found to be under positive selection, every single one of them has been duplicated in at least one lineage of freshwater sponge. These genes are therefore evolving in multiple ways to allow these sponges to survive in freshwater conditions, adding further support to their status as key components of this transition.”

The large number of duplications across freshwater sponge phylogeny is surprising, as their close relationships would suggest that they would contain more shared gene duplications, whereas outgroups would have more gene duplications, accrued over evolutionary time. This is particularly true in the case of Lake Baikal sponges, which are closely related, and diverged from one another relatively recently. This finding is not a consequence of undersampling of genetic resources in our marine outgroups, as three genomes (*A. queenslandica, X. testudinaria* and *T. wilhelma*) were used as comparison points. We hypothesise therefore that freshwater sponges are adjusting to their environment by large-scale duplication of their genes, as well as by changing their sequence under evolutionary pressure as noted earlier. These additional duplicates will provide raw material for evolution in a variety of ways, allowing subfunctionalisation, regulatory changes, and the emergence of novel functions from ancient genes.

To check whether species-specific novelties (sequences not observed in, or assigned to orthogroups shared with, other species), also exhibited this trend, we calculated their abundance as can be seen in Figure 6B. In contrast with our gene duplication results, freshwater sponges exhibited few novel genes when compared to outgroup taxa. Similarly, Fig 6C shows species-unique orthogroups, where more than one sequence exists for a gene, but only in a single species. This shows a similar distribution, with species-specific orthogroups more common in our marine outgroups. These findings will be biased somewhat by the close relationship between freshwater taxa, making it more likely that novelties are shared by these species. However, it is important to note that freshwater taxa, and particularly Lake Baikal sponges, do not exhibit large amounts of novelty on an individual species level.

It is difficult to assign identity or function to these genes, as they are novelties to the individual lineages or clades in which they are found. In the case the novelties noted within the Spongillidae by Orthofinder, they have no shared orthologue within any of the marine outgroups shown in Fig 6A, including three genomic resources, which should be reasonably complete datasets. The genes recorded here are therefore highly likely to be true novelties. They may, however, belong to larger gene families, or possess some sequence signatures which give clues as to their identity. It will be particularly interesting to compare these novelties to those found in freshwater-adapted species from other phyla, when such data become available.

To examine the extent to which orthologues and orthogroups are shared between the species examined here, we plotted this data on a matrix (Figure 6D), and used colour shading to display it as a heat map. It is clear from this data that in general freshwater sponges share more sequences in common than the other outgroup species in our dataset do with one another. Only *M. phyllophilla* and *T. wilhema* are similar to the same extent, and *M. phyllophilla* also seems to share a conserved gene set with most of the other sponges in our sampling. In contrast, *H. tubifera, A. queenslandica* and *X. testudinaria* are the species with the most divergent gene cassettes. This is not a consequence of incomplete or inadequate transcriptomic sampling, as the latter two species are represented by genomic sequence. At the other end of the spectrum is *E. fragilis* and *E. muelleri* which share the highest similarity, and not, as might be expected, our closely related Lake Baikal sponges.

Summarising the evidence presented in Figure 6, freshwater sponges therefore possess a large number of novel genes (Fig. 6A), generated largely through duplication rather than “*de novo*” (Fig. 6B,C). These cassettes are well conserved in freshwater sponges, but Lake Baikal sponges are not the most prominent examples of this (Fig. 6D) – other freshwater sponges have more complete gene sets, with better conservation of orthogroups. However, Lake Baikal sponges possess many duplication-derived novelties of their own (Fig 6A) and these are likely to be the source of many of the unique traits possessed by the Lubomirskidae.

## Conclusions

The animals of Lake Baikal in general, and the freshwater sponges found there in particular, possess a number of unique biological features. Here we have presented data, in the form of three deep transcriptomes and a draft genome, allowing further investigations of the biology of these fascinating organisms. Using these, we have examined the adaptation of these sponges to freshwater environments from a number of angles. A disparate, but vital, list of genes shows strong evidence for positive selection, especially in genes related to membrane function and transport. These sponges have used gene duplication to generate the raw material for evolution, and have leveraged symbioses to allow them to live in challenging conditions. These findings provide a crucial comparison point when establishing the fundamental requirements, at the molecular level, for evolution to freshwater conditions. Lake Baikal sponges have much to teach us about the myriad demands of adaptation to freshwater and to their unique habitat, and this research will be fruitful for many years to come.

## Materials and Methods

### Sponge Collection, Nucleic Acid Extraction, Library Construction and Next Generation Sequencing

Sponge samples from healthy individuals of the three species studied (one specimen per species) were collected by SCUBA diving in September 2015 in the Southern basin of Lake Baikal near the village of Bolshie Koty (west coast, 51°53’54.4“N 105°04’15.3”E) at 10m depth. Samples of tissue for RNA extraction were placed in IntactRNA (Evrogen) immediately on return to the surface, incubated at 4°C overnight, before transfer to a −80°C freezer for long term storage. Samples for DNA extraction were placed in 100% ethanol and stored at −20° C. The *L. baikalensis* sample for genomic study was taken from the same individual as the mRNA sample.

RNA extraction was performed using a ExtractRNA kit (Evrogen). An Agilent Technologies 2100 Bioanalyzer or 2200 TapeStation were used to establish that the RNA Integrity Number (RIN) value was greater than or equal to 8 for samples for sequencing. RNA libraries were constructed by Macrogen using the TruSeq RNA Sample Prep Kit v2. The HiSeq2000 platform (Illumina, USA) was used for paired-end sequencing at 150 bp.

DNA extraction was performed using a standard CTAB method (Gustincich et al 1991). DNA quality was assessed by running a subsample on a 1% agarose gel and the quantity of DNA was measured using a NanoVue (GE Healthcare). DNA libraries were constructed by Macrogen using a TruSeq DNA PCR-free library kit (350bp insert size). An Illumina Hiseq X10 platform was used for sequencing, performed by Macrogen. Initial assessment of read quality and de-multiplexing of reads was performed by the provider according to their proprietary procedures. Paired-end reads were made available for download from their server, with no unpaired orphan reads retained by the process.

### Initial Quality Assessment

The FastQC program (Andrews 2010) was used to perform initial quality assessment of all reads. The RNA reads required some cleaning, which was performed using Trimmomatic 0.33 (Bolger et al 2014) with the following settings: ILLUMINACLIP:../Adaptors.fa:2:30:10 LEADING:3 TRAILING:3 SLIDINGWINDOW:4:20 MINLEN:30. The Adaptors.fa file was adjusted to include the adaptor sequences specific to each read pair. Several problems were observed with DNA reads in particular, as detailed in the results. To remedy these, Trimmomatic was first run as described above, before rCorrector (Song and Florea 2015) was used to correct reads. Read pairs for which one of the reads was unfixable were removed using the FilterUncorrectablePEfasta.py script (https://github.com/harvardinformatics/). FastQC was then re-run to confirm the quality of reads before assembly. Further, seqtk fqchk (https://github.com/lh3/seqtk) was used to ascertain additional basic metrics.

### Assembly and Assessment

Assemblies of RNAseq data were performed using Trinity version 2013_08_14 (Grabherr et al 2011). Samples from *B. bacillifera* (A2) *Lubomirskia abietina* (A10) and *Lubomirskia baikalensis* (A8) were assembled separately. For our genomic data, a variety of assembly methods were trialed at a range of *k* mer sizes. Velvet 1.2.10 (Zerbino and Birney 2008), ABySS 2.0.2 (Simpson et al 2009), SOAPdenovo2 2.04 (Luo et al 2012) and SPAdes 3.9.1 (Bankevich et al 2012) were assayed as noted in the results. Default settings were used with the exception of a minimum contig size of 200 and a minimum coverage of 3 when possible. As these assemblies were to be used for the identification of symbiote and bacterial sequence, no cleaning was performed after assembly.

Numerical metrics relating to assembly were recovered using basic perl and python scripts, available from the authors on request. BUSCO v1.1b1 (Simao 2015) was used to assess the content of both the genome and transcriptomic assemblies, with the completeness of these measured relative to the eukaryotic and metazoan Basic Universal Single Copy Orthologue (BUSCO) cassettes. *K* mers were counted using Jellyfish (Marcais and Kingsford 2011) and GenomeScope (Verture et al 2017) was used to assay the genome size, heterozygosity and coverage of the *L. baikalensis* genome sample.

To assess the content of these assemblies further, Blobtools (Laetsch and Blaxter 2017) was run, using Diamond (Buchfink et al 2015) to run BLASTx searches for assignment of taxon identity, against the NCBI *nr* database (*E* value cutoff: 1e-8, with best hit retained). Bowtie2 (Langmead and Salzberg 2012) was used for read mapping. Blobtools was then run using the *de novo* genome pathway, and with mapping against a local copy of TaxID files.

### Gene Identification and Annotation

For individual genes, tBLASTn (Altschul et al 1990) was used on a locally constructed database to putatively identify genes using sequences of known homology downloaded from Genbank. To further confirm identity, the sequences thus identified were reciprocally BLASTed (BLASTx) against the NCBI nr database online using BLASTx.

For total annotation, the longest ORF for each contig was translated using the getORF.py python script, retaining only the longest ORF, followed by BLASTP annotation (Altschul et al 1990) against the *nr* protein database. These results were imported into Blast2GO Pro (Gotz et al 2008) followed by InterPro scanning, mapping, annotation (including ANNEX augmentation) and enzyme code mapping. The complete annotations for these are attached as Supplementary File 4.

### Phylogenetic Analyses of Symbiont Sequences

The complete 18S rRNA sequence of a potential dinoflagellate symbiont recovered in our genomic sample of *L. baikalensis* (NODE_102849_length_610_cov_109.672) was aligned with 58 other sequences of *Gyrodinium* spp. and uncultured marine and freshwater eukaryotes obtained in NCBI (Supplementary File 5) using Muscle (Edgar 2004) in SeaView v4.0 (Gouy et al. 2009). The phylogenetic analysis was performed using GAMMA + G + I as model of substitution in RAxML v8.0 (Stamatakis 2014) with 10 runs and 100 bootstrap replicates. Also, the sequence of the partial 18S-ITS1 rRNA identified in blast as a Chlorophyta (NODE_291_length_24938_cov_9.23424) was aligned with 57 sequences of other Chlorophyta (including *Choricystis*, *Chlorella*, *Chloroidium*, *Chlorocloster*, *Botryococcus*, *Pseudococcomyxa*, and *Coccomyxa*) obtained in NCBI (Supplementary File 5) using the same alignment strategy as before. Phylogenetic analysis was performed as previously.

### Mitochondrial Sequence Identification, Annotation and Phylogenetic Analysis

TBLASTN (Altschul et al 1990) was used to identify mitochondrial sequences (- db_gencode 4) using sequences of known homology from other sponges. This recovered a complete circular mitochondrial sequence after manual alignment and removal of extraneous sequences. Annotation was performed using the MITOS2 webserver (Bernt et al 2013), with the “04 - Mold/Protozoan/Coelenterate” setting used as the translational code for sponge data. Manual curation was performed to confirm start/stop codon identity, using homology to known genes to collaborate start/stop location. Mitochondrial phylogeny was inferred based on nucleotide sequence of all of these genes (both protein coding and rRNA), alongside those of known homology downloaded from Genbank. These were aligned on a gene-by-gene basis using MAFFT (Katoh et al 2002), and the G-INS-i approach. After concatenation with FASConCat-G (Kuck and Longo 2014) gaps and regions of poor alignment were excluded and phylogenetic inference was performed using partitioned Bayesian analysis in MrBayes (Ronquist and Huelsenbeck 2003), with models chosen using mixed models in the first instance. The first 25% of samples were discarded as ‘burn-in’, before the remaining samples were used to generate the figures shown here. Alignments are displayed in Geneious (Kearse et al 2012) for figure generation.

### Selection Test

Transdecoder (Haas et al 2013), Orthofinder (Emms and Kelly 2018, Diamond, Fasttree, MCL and MAFFT options) and Phylotreepruner (Kocot et al 2013) were run sequentially to identify a dataset of orthologous gene alignments. Final alignments, after pruning, are represented by only one sequence per species (i.e. no paralogous sequence). Concatenated amino acid sequence from these alignments were used as the basis for partitioned Bayesian analysis in MrBayes (Ronquist and Huelsenbeck 2003) to determine a species tree for selection tests (shown in Fig 4A), with a GTR+4G model applied to each gene in an independent partition. Selection tests were performed according to a schema put forward in Santagata (2018). PAL2NAL (Suyama et al 2006) was then run orthogroup-by-orthogroup to find the CDS region corresponding to the aligned protein sequence, and generate a nucleotide alignment for performing selection tests.

CODEML was run in PAML (Muse and Gaut 1994, Yang 2007) to test null vs alternative hypotheses as to gene-level selection, and differences in LnL used as the basis for χ^2^ tests of level of significance. These were corrected for multiple comparison FDR using Benjamini and Hochberg correction (Benjamini and Hochberg 1995, Benjamini and Yekutieli 2001). Bayes Empirical Bayes (BEB) values in the ALT output were also extracted and used to identify sites under selection (Yang et al 2005). HyPHy was run, with BUSTED, aBSREL and MEME tests of branch-level and site-level selection respectively (Pond et al 2005, Murrell et al 2012, Murrell et al 2015, Smith et al 2015). Sequences were annotated by BLASTP identity to the nr database.

### GO Over-representation analysis

To identify GO terms over-represented in the gene set noted as under selection by all tests, we used the ShinyGO (Ge and Jung 2018) tool. The accession numbers of the *A. queenslandica* orthologues within the positively selected orthogroups were input into ShinyGo, with a P-value cutoff for FDR of 0.05. ShinyGO automatically retrieved the GO terms for these genes from the annotated *A. queenslandica* resource, determined enriched GO terms within this set and corrected for FDR. The results of over-representation analyses, alongside network diagrams, were downloaded from the ShinyGO server. To visualise over-enriched GO complements, the Revigo tool (Supek et al 2011) was used to display this data, with these settings: List: Medium 0.7, database: whole Uniprot, Semantic similarity: Simrel. This was exported as an R plot, and a pdf generated. Font and colour was edited in Adobe Illustrator (Adobe: San Jose, California) for display.

### Novelty Identification

To identify novelties shared across freshwater sponges, but absent from outgroups, we used the same datasets as used for the selection tests described above, with the addition of the protein sets from the genomes of *Xestospongia testudinaria* (Ryu et al 2016) and *Tethya wilhelmi* (Francis et al 2017) to compare with complete gene sets as outgroups. Orthofinder2 (Emms and Kelly 2018) was run to identify orthogroups (settings: Diamond, Fasttree, MCL and MAFFT), and to identify those found uniformly in freshwater datasets, but not in outgroups. This data was used to extract data on gene duplication at nodes, statistics regarding numbers of genes assigned to orthogroups, and shared orthogroup and orthologue figures, as displayed in Fig 6.

## Supplementary Files

*Supplementary File 1:* Further details on sequencing and assembly of resources. Tables containing statistics related to reads, assemblies and gene complements as assessed with BUSCO. (.pdf)

*Supplementary File 2:* GenomeScope profile of *k-*mer distribution, based on Jellyfish *k-*mer counts of untrimmed and uncorrected reads. Note peak of *k-*mer coverage (15.2) and inferred genome size of 565,078,853 bp. (.pdf)

*Supplementary File 3:* Mapping results for transcriptomic and genome assemblies vs. the nr database. %ages represent proportion of base pairs. At left: overall %age of contigs mapped to a known sequence. At right, Superkingdom and (inset) Phylum of these hits. Top 6 (Superkingdom)/Top 7 (Phylum) results shown. (.pdf)

*Supplementary File 4:* Annotations for transcriptomes (.annot, zipped). Available from Figshare https://figshare.com/s/fe36239c32bbf7342756 doi: <u>10.6084/m9.figshare.6819812</u> *Supplementary File 5:* Details of Blobplot taxonomic binning, hits and alignments of sequences with best BLAST identity to *Symbiodinium microadriaticum* (.xlsx, .txt files). Accession numbers, alignments and sequences of symbiont sequences used in phylogenetic analysis (.phy, .fa, .xlsx files). Details of selection tests, and complete site-by-site data for significant genes (.xlsx files). GO assignation, REVIGO data and scripts used to create Fig 5. All in single .zip file. Available from Figshare https://figshare.com/s/fe36239c32bbf7342756 doi: <u>10.6084/m9.figshare.6819812</u>

## Data Note

Raw reads are available from the NCBI SRA under accession PRJNA431612. Assemblies, including preliminary and alternate forms, have been uploaded to Figshare, with DOI: and URL: 10.6084/m9.figshare.6819812, https://figshare.com/s/fe36239c32bbf7342756.

## Contribution of Authors

VBI conceived of the study, gathered specimens and performed RNA and DNA extractions. NJK, BP, AR and VBI designed and performed experiments. BP performed mitochondrial data analysis. NJK performed assembly and bioinformatic analysis. All authors wrote and agreed on the final form of the manuscript.

## Acknowledgements

This work was supported by the ADAPTOMICS MSCA [grant number: IF750937] with a grant to NJK under the Horizon 2020 program. This study was done in the framework of the State project 0345-2019-0002 (АААА-А16-116122110066-1) and supported by RFBR grants No 17-04-01598, 17-44-388103 (genome and transcriptome sequencing) to VBI. Many of the selection test methods applied in this study were facilitated using best practices and scripts provided as part of a Next Generation Sequencing-based workshop sponsored by the National Science Foundation (Award # 1744877 to S. Santagata) and we are thank Dr Santagata for his guidance with this protocol. The authors thank the members of their laboratories for their many helpful discussions and support. We thank editors and two anonymous reviewers for their time and thoughtful contributions towards this manuscript.

## References

Altschul SF, Gish W, Miller W, Myers EW, Lipman DJ. 1990. Basic local alignment search tool. J Mol Biol. 215:403–410.

Alverson AJ. 2007. Strong purifying selection in the silicon transporters of marine and freshwater diatoms. Limnol Oceanog. 52:1420–9.

Andrews S. 2010. FastQC: A quality control tool for high throughput sequence data [Internet]. http://www.bioinformatics.babraham.ac.uk/projects/fastqc/

Annenkova NV, Lavrov DV, Belikov SI. 2011. Dinoflagellates associated with freshwater sponges from the ancient Lake Baikal. Protist. 162:222–36.

Bankevich A, Nurk S, Antipov D, Gurevich AA, Dvorkin M, Kulikov AS, Lesin VM, Nikolenko SI, Pham S, Prjibelski AD, Pyshkin AV. 2012. SPAdes: a new genome assembly algorithm and its applications to single-cell sequencing. J Comp Biol. 19:455–477.

Benjamini Y, Hochberg Y. 1995. Controlling the false discovery rate: a practical and powerful approach to multiple testing. J Roy Stat Soc B. 1:289–300.

Benjamini Y, Yekutieli D. 2001. The control of the false discovery rate in multiple testing under dependency. Ann Stat. 1:1165–1188.

Bernt M, Donath A, Jühling F, Externbrink F, Florentz C, Fritzsch G, Pütz J, Middendorf M, Stadler PF. 2013. MITOS: Improved de novo metazoan mitochondrial genome annotation. Mol Phylog Evol. 69:313–319.

Bielawski JP, Baker JL, Mingrone J. 2016. Inference of Episodic Changes in Natural Selection Acting on Protein Coding Sequences via CODEML. Curr Protoc Bioinformatics. 54:1–32.

Bolger AM, Lohse M, Usadel B. 2014. Trimmomatic: a flexible trimmer for Illumina sequence data. Bioinformatics. 30:2114–2120.

Buchfink B, Xie C, Huson, DH. 2015. Fast and sensitive protein alignment using DIAMOND. Nat Methods. 12:59.

Bustelo XR, Sauzeau V, Berenjeno IM. 2007. GTP binding proteins of the Rho/Rac family: regulation, effectors and functions in vivo. Bioessays. 29:356–370.

Cárdenas P, Pérez T, Boury-Esnault N. 2012. Sponge Systematics facing new challenges. Adv Mar Biol. 61:79–209.

Chernogor L, Denikina N, Kondratov I, Solovarov I, Khanaev I, Belikov S, Ehrlich H. 2013. Isolation and identification of the microalgal symbiont from primmorphs of the endemic freshwater sponge *Lubomirskia baicalensis* (Lubomirskiidae, Porifera). Eur J Phycol. 48:497–508.

Ciesielski TM, Pastukhov MV, Leeves SA, Farkas J, Lierhagen S, Poletaeva VI, Jenssen BM. 2016. Differential bioaccumulation of potentially toxic elements in benthic and pelagic food chains in Lake Baikal. Env Sci Poll Res. 23:15593–15604.

Costa R, Keller-Costa T, Gomes NC, da Rocha UN, van Overbeek L, van Elsas JD. 2013. Evidence for selective bacterial community structuring in the freshwater sponge Ephydatia fluviatilis. Microbial ecology. 65:232–44.

Darriba D, Taboada GL, Doallo R, Posada D. 2011. ProtTest 3: fast selection of best-fit models of protein evolution. Bioinformatics. 27:1164–1165.

DeFaveri J, Shikano T, Shimada Y, Goto A, Merilä J. 2011. Global analysis of genes involved in freshwater adaptation in threespine sticklebacks (*Gasterosteus aculeatus*). Evolution. 65:1800–1807.

Denikina NN, Dzyuba EV, Bel’kova NL, Khanaev IV, Feranchuk SI, Makarov MM, Granin NG, Belikov SI. 2016. The first case of disease of the sponge *Lubomirskia baicalensis:* Investigation of its microbiome. Biol Bull. 43:263–270.

Edgar RC. 2004. MUSCLE: multiple sequence alignment with high accuracy and high throughput. Nucl acids res, 32:1792–1797.

Emms DM, Kelly S. 2018. OrthoFinder2: fast and accurate phylogenomic orthology analysis from gene sequences. BioRxiv. 1:466201.

Erpenbeck D, Weier T, De Voogd NJ, Wörheide G, Sutcliffe P, Todd JA, Michel E. 2011. Insights into the evolution of freshwater sponges (Porifera: Demospongiae: Spongillina): barcoding and phylogenetic data from Lake Tanganyika endemics indicate multiple invasions and unsettle existing taxonomy. Mol Phylog Evol. 61:231–6.

Excoffier L, Lischer HE. 2010. Arlequin suite ver 3.5: a new series of programs to perform population genetics analyses under Linux and Windows. Mol Ecol Res. 10:564–567.

Feranchuk S, Belkova N, Chernogor L, Potapova U, Belikov S. 2018. The signs of adaptive mutations identified in the chloroplast genome of the algae endosymbiont of Baikal sponge. F1000Research. 4:7.

Fernandez-Valverde SL, Calcino AD, Degnan BM. 2015. Deep developmental transcriptome sequencing uncovers numerous new genes and enhances gene annotation in the sponge *Amphimedon queenslandica*. BMC Genomics. 16:387.

Francis WR, Eitel M, Vargas S, Adamski M, Haddock SH, Krebs S, Blum H, Erpenbeck D, Wörheide G. 2017. The Genome Of The Contractile Demosponge *Tethya wilhelma* And The Evolution Of Metazoan Neural Signalling Pathways. bioRxiv. 1:120998.

Gaikwad S, Shouche YS, Gade WN. 2016. Microbial community structure of two freshwater sponges using Illumina MiSeq sequencing revealed high microbial diversity. AMB Express. 6:40.

Ge S, Jung D. 2018. ShinyGO: a graphical enrichment tool for animals and plants. bioRxiv. 1:315150.

Gernert C, Glöckner FO, Krohne G, Hentschel U. 2005. Microbial diversity of the freshwater sponge *Spongilla lacustris*. Microbial ecology. 50:206–12.

Goodwin S, McPherson JD, McCombie WR. 2016. Coming of age: ten years of next-generation sequencing technologies. Nat Rev Genetics. 17:333–351.

Goldman N, Yang Z. 1994. A codon-based model of nucleotide substitution for protein-coding DNA sequences. Mol Biol Evol. 11:725–736.

Götz S, García-Gómez JM, Terol J, Williams TD, Nagaraj SH, Nueda MJ, Robles M, Talón M, Dopazo J, Conesa A. 2008. High-throughput functional annotation and data mining with the Blast2GO suite. Nucleic Acids Res. 36:3420–35.

Gouy M, Guindon S, Gascuel O. 2009. SeaView version 4: a multiplatform graphical user interface for sequence alignment and phylogenetic tree building. Mol Biol Evol. 27:221–224.

Grabherr MG, Haas BJ, Yassour M, Levin JZ, Thompson DA, Amit I, et al. 2011. Full-length transcriptome assembly from RNA-Seq data without a reference genome. Nat Biotechnol. 29:644–652.

Gupta V, Bamezai RN. 2010. Human pyruvate kinase M2: a multifunctional protein. Prot Sci. 19:2031–44.

Gustincich S, Manfioletti G, Del GS, Schneider C, Carninci P. 1991. A fast method for high-quality genomic DNA extraction from whole human blood. Biotechniques. 11:298–300.

Hansen KD, Brenner SE, Dudoit S. 2010. Biases in Illumina transcriptome sequencing caused by random hexamer priming. Nucleic Acids Res. 38:e131.

Haas BJ, Papanicolaou A, Yassour M, Grabherr M, Blood PD, Bowden J, Couger MB, Eccles D, Li B, Lieber M, MacManes MD. 2013. De novo transcript sequence reconstruction from RNA-seq using the Trinity platform for reference generation and analysis. Nature Protoc. 8:1494.

Itskovich VB, Kaluzhnaya OV, Veynberg E, Erpenbeck DI. 2015. Endemic Lake Baikal sponges from deep water. 1: Potential cryptic speciation and discovery of living species known only from fossils. Zootaxa. 3990:123–137.

Itskovich VB, Shigarova AM, Glyzina OY, Kaluzhnaya OV, Borovskii GB. 2018. Heat shock protein 70 (Hsp70) response to elevated temperatures in the endemic Baikal sponge *Lubomirskia baicalensis*. Ecological Indicators. 88:1–7.

Jeffery NW, Jardine CB, Gregory TR. 2013. A first exploration of genome size diversity in sponges. Genome. 56:451–456

Kaluzhnaya O, Itskovich VB, McCormack GP. 2011. Phylogenetic diversity of bacteria associated with the endemic freshwater sponge *Lubomirskia baicalensis*. World J Microbiol Biotechnol. 27:1955–1959.

Kaluzhnaya OV, Krivich AA, Itskovich VB. 2012. Diversity of 16S rRNA genes in metagenomic community of the freshwater sponge *Lubomirskia baicalensis*. Russ J Genet 48:855–858.

Kaluzhnaya OV, Itskovich VB. 2015. Bleaching of Baikalian sponge affects the taxonomic composition of symbiotic microorganisms. Russ J Genet. 51:1153–1157.

Kaluzhnaya OV, Itskovich VB. 2017. Molecular identification of filamentous cyanobacteria overgrowing the endemic sponge *Lubomirskia baicalensis*. Inland Waters. 7:267–271.

Kasimov N, Karthe D, Chalov S. 2017. Environmental change in the Selenga River-Lake Baikal Basin. Regional Environmental Change. 17:1945–1949.

Katoh K, Misawa K, Kuma KI, Miyata T. 2002. MAFFT: a novel method for rapid multiple sequence alignment based on fast Fourier transform. Nucleic Acids Res. 30:3059–3066.

Kearse M, Moir R, Wilson A, Stones-Havas S, Cheung M, Sturrock S, Buxton S, Cooper A, Markowitz S, Duran C, Thierer T. 2012. Geneious Basic: an integrated and extendable desktop software platform for the organization and analysis of sequence data.Bioinformatics. 28:1647–1649.

Kenny NJ, de Goeij JM, de Bakker DM, Whalen CG, Berezikov E, Riesgo A. 2018. Towards the identification of ancestrally shared regenerative mechanisms across the Metazoa: A Transcriptomic case study in the Demosponge *Halisarca caerulea*. Marine Genomics. 37:135–47.

Khanaev IV, Kravtsova LS, Maikova OO, Bukshuk NA, Sakirko MV, Kulakova NV, Butina TV, Nebesnykh IA, Belikov SI. 2018. Current state of the sponge fauna (Porifera: Lubomirskiidae) of Lake Baikal: Sponge disease and the problem of conservation of diversity. Journal of Great Lakes Research. 44:77–85.

Kocot KM, Citarella MR, Moroz LL, Halanych KM. 2013. PhyloTreePruner: a phylogenetic tree-based approach for selection of orthologous sequences for phylogenomics. Evolut Bioinform. 9:EBO-S12813.

Kozhov M. 1963. Lake Baikal and its life, vol. XI of. Monographia Biologicae. Dr. W. Junk Publishers: The Hague.

Kozhova OM, Izmest’eva LR. (Eds). 1998. Lake Baikal – Evolution and Biodiversity Backhuys Publishers: Leiden.

Krienitz L, Huss VAR, Hümmer C. 1996. Picoplanktonic *Choricystis* species (Chlorococcales, Chlorophyta) and problems surrounding the morphologically similar *’Nannochloris*-like algae’. Phycologia. 35:332–341.

Kulakova NV, Sakirko MV, Adelshin RV, Khanaev IV, Nebesnykh IA, Perez T. 2017. Brown Rot Syndrome and Changes in the Bacterial Сommunity of the Baikal Sponge *Lubomirskia baicalensis*. Microb Ecol. 1:1–11.

Kück P, Longo GC. 2014. FASconCAT-G: extensive functions for multiple sequence alignment preparations concerning phylogenetic studies. Front Zool. 11:81.

Laetsch DR, Blaxter ML. 2017. BlobTools: Interrogation of genome assemblies. F1000Research. 6:1.

Langmead B, Salzberg SL. 2012. Fast gapped-read alignment with Bowtie 2. Nat Methods. 9:357–359.

Latyshev NA, Zhukova NV, Efremova SM, Imbs AB, Glysina OI. 1992. Effect of habitat on participation of symbionts in formation of the fatty acid pool of fresh-water sponges of Lake Baikal. Comp Biochem Physiol B: Comp Biochem. 102:961–5.

Lavrov DV. Maikova OO, Pett W, Belikov SI. 2012. Small inverted repeats drive mitochondrial genome evolution in Lake Baikal sponges. Gene. 505:91–99.

Li B, Dewey CN. 2011. RSEM: accurate transcript quantification from RNA-Seq data with or without a reference genome. BMC Bioinformat. 12:323.

Liu F, Li J, Feng G, Li Z. 2016. New genomic insights into “Entotheonella” symbionts in *Theonella swinhoei*: Mixotrophy, anaerobic adaptation, resilience, and interaction. Front Microbiol. 25:1333.

Lohse M, Drechsel O, Bock, R. 2007. OrganellarGenomeDRAW (OGDRAW): a tool for the easy generation of high-quality custom graphical maps of plastid and mitochondrial genomes. Curr Genet. 52:267–274.

Luo R, Liu B, Xie Y, Li Z, Huang W, Yuan J, He G, Chen Y, Pan Q, Liu Y, Tang J. 2012. SOAPdenovo2: an empirically improved memory-efficient short-read de novo assembler. GigaScience. 1:1–18.

Mäkinen HS, Cano JM, Merilä J. 2008. Identifying footprints of directional and balancing selection in marine and freshwater three spined stickleback (*Gasterosteus aculeatus*) populations. Mol Ecol. 17:3565–3582.

Marçais G, Kingsford C. 2011. A fast, lock-free approach for efficient parallel counting of occurrences of k-mers. Bioinformatics. 27:764–770.

Manconi R. Pronzato R. 2002. Suborder Spongillina subord. nov.: Freshwater sponges. 921-1020. In Hooper JNA, Van Soest RWM. (ed.) Systema Porifera. A guide to the classification of sponges 1. Kluwer Academic/Plenum Publishers: New York.

Manconi R, Pronzato R. 2008. Global diversity of sponges (Porifera: Spongillina) in freshwater. Hydrobiologia. 595:27–33.

Meixner MJ, Lüter C, Eckert C, Itskovich V, Janussen D, von Rintelen T, Bohne AV, Meixner JM, Hess WR. 2007. Phylogenetic analysis of freshwater sponges provide evidence for endemism and radiation in ancient lakes. Mol Phylogenet Evol. 45:875–886.

Morrow C, Cárdenas P. 2015. Proposal for a revised classification of the Demospongiae (Porifera). Front Zool. 12:7.

Müller WE, Belikov SI, Kaluzhnaya OV, Perović Ottstadt S, Fattorusso E, Ushijima H, Krasko A, Schröder HC. 2007. Cold stress defense in the freshwater sponge *Lubomirskia baicalensis*. The FEBS journal. 274:23–36.

Müller WE, Belikov SI, Kaluzhnaya OV, Chernogor L, Krasko A, Schröder HC. 2009. Symbiotic interaction between dinoflagellates and the demosponge *Lubomirskia baicalensis*: aquaporin-mediated glycerol transport. In Müller WE, Grachev MA. (ed.) Biosilica in evolution, morphogenesis, and nanobiotechnology. Springer, Berlin, Heidelberg.

Murrell B, Wertheim JO, Moola S, Weighill T, Scheffler K, Kosakovsky Pond SL. 2012. Detecting Individual Sites Subject to Episodic Diversifying Selection. PLoS Genet. 8:e1002764.

Murrell B, Weaver S, Smith MD, Wertheim JO, Murrell S, Aylward A, Eren K, Pollner T, Martin DP, Smith DM, Scheffler K, Kosakovsky Pond SL. 2015. Gene-Wide Identification of Episodic Selection. Mol Biol Evol. 32:1365–1371.

Muse SV, Gaut BS. 1994. A likelihood approach for comparing synonymous and nonsynonymous nucleotide substitution rates, with application to the chloroplast genome. Mol Biol Evol. 11:715–724.

Narum SR. 2006. Beyond Bonferroni: less conservative analyses for conservation genetics. Cons Genet. 7:783–787.

Naumenko SA, Logacheva MD, Popova NV, Klepikova AV, Penin AA, Bazykin GA, Etingova AE, Mugue NS, Kondrashov AS, Yampolsky LY. 2017. Transcriptome based phylogeny of endemic Lake Baikal amphipod species flock: fast speciation accompanied by frequent episodes of positive selection. Mol Ecol. 26:536–553.

Pile AJ, Patterson MR, Savarese M, Chernykh VI, Fialkov VA. 1997. Trophic effects of sponge feeding within Lake Baikal’s littoral zone: 2. Sponge abundance, diet, feeding efficiency, and carbon flux, Limno Oceonogr. 42:178–184.

Pomazkina G, Kravtsova L, Sorokovikova E. 2012. Structure of epiphyton communities on Lake Baikal submerged macrophytes. Limno Rev. 12:19–27.

Pond SLK, Frost SDW, Muse SV. 2005. HyPhy: hypothesis testing using phylogenies. Bioinformatics. 21:676–679.

Rahi ML, Amin S, Mather PB, Hurwood DA. 2017. Candidate genes that have facilitated freshwater adaptation by palaemonid prawns in the genus *Macrobrachium*: identification and expression validation in a model species (*M. koombooloomba*). PeerJ. 5:e2977.

Rice P, Longden I, Bleasby A, 2000. EMBOSS: the European molecular biology open software suite. Trends Genet. 16:276–7.

Riesgo A, Farrar N, Windsor PJ, Giribet G, Leys SP. 2014. The analysis of eight transcriptomes from all poriferan classes reveals surprising genetic complexity in sponges. Mol Biol Evol. 31:1102–1120.

Rivarola-Duarte L, Otto C, Jühling F, Schreiber S, Bedulina D, Jakob L, Gurkov A, Axenov-Gribanov D, Sahyoun AH, Lucassen M, Hackermüller J. 2011. A first glimpse at the genome of the Baikalian amphipod Eulimnogammarus verrucosus. J Exp Biol B. 322:177–189.

Robinson MD, McCarthy DJ, Smyth GK. 2010. edgeR: a Bioconductor package for differential expression analysis of digital gene expression data. Bioinformatics. 26:139–40.

Romanova EV, Kravtsova LS, Izhboldina LA, Khanaev IV, Sherbakov DY. 2015. Identification of filamentous green algae from an area of local biogenic pollution of Lake Baikal (Listvennichnyi Bay) using SSU 18S rDNA. Russ J Genet: App Res. 5:118–125.

Romanova EV, Aleoshin VV, Kamaltynov RM, Mikhailov KV, Logacheva MD, Sirotinina EA, Gornov AY, Anikin AS, Sherbakov DY. 2016. Evolution of mitochondrial genomes in Baikalian amphipods. BMC Genomics. 17:1016.

Ronquist F, Huelsenbeck JP, 2003. MrBayes 3: Bayesian phylogenetic inference under mixed models. Bioinformatics. 19:1572–1574.

Rusinek OT, Takhteev VV, Gladkochub DP, Khodzher TV, Budnev NM, editors. 2012a. Baicalogy. Book 1. Nauka Publishers: Novosibirsk.

Rusinek OT, Takhteev VV, Gladkochub DP, Khodzher TV, Budnev NM, editors. 2012b. Baicalogy. Book 2. Nauka Publishers: Novosibirsk.

Ryu T, Seridi L, Moitinho-Silva L, Oates M, Liew YJ, Mavromatis C, Wang X, Haywood A, Lafi FF, Kupresanin M, Sougrat R. 2016. Hologenome analysis of two marine sponges with different microbiomes. BMC Genomics. 17:158.

Sand-Jensen K, Pedersen MF. 1994. Photosynthesis by symbiotic algae in the freshwater sponge, Spongilla lacustris. Limno Oceanog. 39:551–61.

Santagata S, 2018, Polar Workshop Workflow Github repository, Github, https://github.com/Santagata/Select_Test,

Sasikumar AN, Perez WB, Kinzy TG. 2012. The many roles of the eukaryotic elongation factor 1 complex. Wiley Interdisciplinary Reviews: RNA. 3:543–55.

Schröder HC, Efremova SM, Itskovich VB, Belikov S, Masuda Y, Krasko A, Müller IM, Müller WE. 2003. Molecular phylogeny of the freshwater sponges in Lake Baikal. J Zool Sys Evol Res. 41:80–86.

Schuster A, Vargas S, Knapp IS, Pomponi SA, Toonen RJ, Erpenbeck D, Wörheide G. 2018. Divergence times in demosponges (Porifera): first insights from new mitogenomes and the inclusion of fossils in a birth-death clock model. BMC Evol Biol. 18:114.

Seo EY, Jung D, Belykh OI, Bukshuk NA, Parfenova VV, Joung Y, Kim IC, Yim JH, Ahn TS. 2016. Comparison of bacterial diversity and species composition in three endemic Baikalian sponges. Ann Limnol-Int J Lim. 52: 27–32.

Shimaraev MN, Verbolov VI, Granin NG, Sherstyankin PP, 1994. Physical Limnology of Lake Baikal: A Review. BICER Publishers: Irkutsk and Okayama.

Simao FA, Waterhouse RM, Ioannidis P, Kriventseva E V, Zdobnov EM. 2015. BUSCO: assessing genome assembly and annotation completeness with single-copy orthologs. Bioinformatics. 31:3210–3212.

Simpson JT, Wong K, Jackman SD, Schein JE, Jones SJ, Birol I. 2009. ABySS: a parallel assembler for short read sequence data. Genome Res. 19:1117–1123.

Smith MD, Wertheim JO, Weaver S, Murrell B, Scheffler K, Kosakovsky Pond SL. 2015. Less is more: an adaptive branch-site random effects model for efficient detection of episodic diversifying selection. Mol Biol Evol. 32:1342–1353.

Song L, Florea L. 2015. Rcorrector: efficient and accurate error correction for Illumina RNA-seq reads. GigaScience. 4:48.

Srivastava M, Simakov O, Chapman J, Fahey B, Gauthier ME, Mitros T, Richards GS, Conaco C, Dacre M, Hellsten U, Larroux C. 2010. The *Amphimedon queenslandica* genome and the evolution of animal complexity. Nature. 466:720–726.

Stamatakis A. 2014. RAxML version 8: a tool for phylogenetic analysis and post-analysis of large phylogenies. Bioinformatics. 30:1312–1313.

Supek F, Bošnjak M, Škunca N, Šmuc T. 2011. REVIGO summarizes and visualizes long lists of gene ontology terms. PloS One. 6:e21800.

Suyama M, Torrents D, Bork P. 2006. PAL2NAL: robust conversion of protein sequence alignments into the corresponding codon alignments. Nucleic Acids Res. 34:W609–12.

Tekedar HC, Karsi A, Reddy JS, Nho SW, Kalindamar S, Lawrence ML. 2017. Comparative genomics and transcriptional analysis of *Flavobacterium columnare* strain ATCC 49512. Front Microbiol. 19:588.

Thomas GWC, Hahn MW, Hahn Y. 2017. The effects of increasing the number of taxa on inferences of molecular convergence. Gen Biol Evol. 9:213–221.

Timoshkin OA. 2001. Lake Baikal: fauna diversity, problems of its immiscibility and origin, ecology and exotic communities. In: Timoshkin OA, editor. Index of animal species inhabiting Lake Baikal and its catchment area. Nauka Publishers: Novosibirsk.

Timofeyev MA. 2010. Ecological and physiological aspects of adaptation to abiotic environmental factors in endemic Baikal and Palearctic amphipods. Tomsk State University: Tomsk.

Untergasser A, Cutcutache I, Koressaar T, Ye J, Faircloth BC, Remm M, Rozen SG. 2012. Primer3—new capabilities and interfaces. Nucleic Acids Res. 40:e115.

Venkat A, Hahn MW, Thornton JW. 2018. Multinucleotide mutations cause false inferences of lineage-specific positive selection. Nat Ecol Evol. 2:1280.

Vurture GW, Sedlazeck FJ, Nattestad M, Underwood CJ, Fang H, Gurtowski J, Schatz MC. 2017. GenomeScope: Fast reference-free genome profiling from short reads. Bioinformatics. 1:btx153.

Wilson MC, Mori T, Rückert C, Uria AR, Helf MJ, Takada K, Gernert C, Steffens UA, Heycke N, Schmitt S, Rinke C. 2014. An environmental bacterial taxon with a large and distinct metabolic repertoire. Nature. 506:58.

Yang Z. 2007. PAML 4: Phylogenetic Analysis by Maximum Likelihood. Mol Biol Evol. 24:1586–1591.

Yang Z, Bielawski JP. 2000. Statistical methods for detecting molecular adaptation. Trends Ecol Evol. 15:496–503.

Yang Z, Wong WSW, Nielsen R. 2005. Bayes empirical bayes inference of amino acid sites under positive selection. Mol Biol Evol. 22:1107–1118.

Zerbino DR, Birney E, 2008. Velvet: algorithms for de novo short read assembly using de Bruijn graphs. Genome Res. 18:821–829.

Zhang J. 2005. Evaluation of an Improved Branch-Site Likelihood Method for Detecting Positive Selection at the Molecular Level. Mol Biol Evol. 22:2472–2479.

