## Supplementary File 1 for "Symbiosis, Selection and Novelty: Freshwater Adaptation in the Unique Sponges of Lake Baikal"

**Full Details, Sequencing and Read Cleaning:**

Our transcriptomic sequencing was of reasonably good quality, with *B. bacillifera* (A2) and *L. abietina* (A10) of good average quality (Phred median >30) through to the final base in both read directions. The *L. baikalensis* (A8) sample was of less robust quality, possessing reads which declined in base call confidence score towards the end of the read. The use of Trimmomatic fixed these issues, with few reads removed from samples A2 and A10, while sample A8 had 40.84% of read pairs removed, either due to poor quality or short read length after truncation. Where possible, single unpaired reads were retained from these filtered read pairs. Unpaired reads were used in assembly, but not in read mapping.

Our genomic sequencing required attention before use in genome assembly, due to a large number of apparent singleton *k*-mers (Supplementary File 2), which could be indicative of read errors. Trimmomatic was applied to these reads first, removing and truncating large quantities of reads, leaving just over half of the original read pairs and bases. The 99,040,746 (53.68%) of pairs remaining after Trimmomatic were fed into rCorrector, which corrected 67,842,587 bases in 34,392,948 read pairs. rCorrector also discarded 17,998,543 pairs of reads, which were assessed as being unfixable. This left a total of 81,042,203 read pairs for further analyses. This process is conservative and highly stringent. In a more complete genome assembly, low complexity sequences could be spanned using long reads and a variety of library sizes, but for our purposes, this ensured the quality of sequence data used in assembly, and thus the reliability of assembled data.

**Additional Details, Assembly:**

SPAdes was chosen as the best-performing assembler, as noted in the main text. For our purposes, the higher median contig size, larger number of long (>1kb) contigs and higher number of bases in these 1kb+ contigs (155,282,664 bp) meant that an increased number of long contigs was available for our work. That SPAdes outperformed other assemblers was not in itself a surprise, as its more recent publication and paired assembly graph approach meant that it incorporates methods not always found in other assemblers. However, other assemblers were better performers by some metrics (for example, ABySS generated the largest single contig). The addition of long read data or multiple library sizes would allow us to further scaffold this data in the future.

The number of bases recovered by our SPAdes assembly (209,989,122 at 500 bp minimum contig size) falls short of the estimated genome size of this sponge by a considerable margin, but would comfortably contain the coding sequence of the average sponge genome. It is likely that repetitive sequences are poorly represented in our dataset, due to the single small fragment size used in sequencing.

**Additional Details, Annotation:**

For all three species present in our sampling, the most commonly top-hit species by BLAST searches was the sponge *Amphimedon queenslandica* with 11,328, 12,236 and 12,358 hits respectively. The affinity of this species, the first sequenced poriferan, with our samples is obvious. As *A. queenslandica* is also a demosponge, and has its genome on the *nr* database, it is unsurprising that it is the species with the highest number of BLAST top-hits. The top five most hit species for all three samples also included *Orbicella faveolata, Stylophora pistillata, Exaiptasia pallida* and *Branchiostoma belcheri,* although the order in which these occurred slightly changes from sample to sample. The first three of these species are cnidarians, and *B. belcheri* is a cephalochordate. These species have a relatively slow rate of molecular evolution, likely resulting in their similarity to our sequences under BLAST search.

### Tables

| *Prior to Cleaning* | ***Baikalospongia bacillifera* (A2)** | ***Lubomirskia abietina* (A10)** | ***Lubomirskia baikalensis* (A8)** | ***Lubomirskia baikalensis* gDNA Reads** |
|---|---|---|---|---|
| **Number of Read Pairs** | 50,694,150 | 52,345,337 | 54,439,423 | 184,491,682 |
| **Read Length** | 101 | 101 | 101 | 151 |
| **GC%** | 49 | 49 | 49 | 43.5 |
| **Average Quality** | 34.4 | 34.39 | 33.74 | 37.5 |
| **Total bases** | 10,240,218,300 | 10,573,758,074 | 10,996,763,446 | 55,716,487,964 |
| ***After Cleaning*** | **A2** | **A10** | **A8** | **gDNA Reads** |
| **Number of Read Pairs** | 48,445,101 | 50,007,487 | 32,204,065 | 81,042,203 |
| **GC% (Paired)** | 49 | 49 | 49 | 41 |
| **Average Quality (Paired)** | 36.25 | 36.25 | 36.15 | 39.75 |
| **Total Bases (Paired)** | 9,518,960,084 | 9,822,304,994 | 6,321,981,078 | 19,426,404,098 |
| **Unpaired Reads** | 2129241 | 2214127 | 1381971 | n/a |
| **GC% (Unpaired )** | 50 | 50 | 50 | n/a |
| **Average Quality (Unpaired)** | 33 | 33 | 32.9 | n/a |
| **Total bases (Unpaired)** | 183279397 | 190482063 | 117844996 | n/a |

*Table 1:* Metrics relating to reads, before and after read cleaning.

| | *Baikalospongia bacillifera* (A2) | *Lubomirskia abietina* (A10) | *Lubomirskia baikalensis* (A8) |
|---|---|---|---|
| **Number of Trinity Transcripts** | 80,925 | 93,404 | 81,951 |
| **Number of Trinity 'Genes'** | 54,606 | 62,809 | 54,913 |
| **Min contig length:** | 201 | 201 | 201 |
| **Max contig length:** | 16,639 | 30,430 | 11,157 |
| **Mean contig length:** | 850.38 | 849.74 | 854.37 |
| **N50 contig length:** | 1,595 | 1,628 | 1,572 |
| **Number of contigs >=1kb:** | 20,558 | 22,946 | 21,595 |
| **Number of contigs in N50:** | 12,300 | 13,341 | 12,943 |
| **Number of bases in all contigs:** | 68,817,041 | 79,368,987 | 70,016,550 |
| **Number of bases in contigs >=1kb:** | 44,913,440 | 52,015,444 | 45,916,970 |
| **GC Content of contigs: (%)** | 46.75 | 46.62 | 46.78 |

*Table 2:* Statistics relating to transcriptome assemblies

| | SPAdes, 500 min | SPAdes, 200 min | ABySS, 61mer | SOAPdenovo, 61mer | Velvet, 61mer |
|---|---|---|---|---|---|
| **Min contig length:** | 500 | 200 | 200 | 200 | 200 |
| **Max contig length:** | 124,926 | 124,926 | 216,201 | 99,521 | 21,493 |
| **Mean contig length:** | 1553.28 | 681.89 | 729.9 | 544.51 | 464.06 |
| **Median contig length:** | 845 | 351 | 294 | 324 | 292 |
| **N50 contig length:** | 2,213 | 1,019 | 1,931 | 661 | 544 |
| **Number of contigs:** | 135,191 | 451,479 | 235,631 | 579,486 | 357,804 |
| **Number of contigs >=1kb:** | 54,728 | 54,728 | 22,065 | 49,540 | 28,260 |
| **Number of contigs in N50:** | 19,573 | 53,387 | 14,759 | 95,954 | 72,346 |
| **Number of bases in all contigs:** | 209,989,122 | 307,857,163 | 171,988,112 | 315,536,346 | 166,041,199 |
| **Number of bases in contigs >=1kb:** | 155,282,664 | 155,282,664 | 96,787,949 | 120,598,815 | 51,363,918 |
| **GC Content of contigs: (%)** | 43.68 | 42.64 | 43.85 | 40.94 | 44.39 |

*Table 3:* Genome assembly using a variety of programmes. SPAdes (with a minimum size cutoff of 500 bp) used for further analyses, but other assemblies also available for download.

| **property** | **min** | **max** |
|---|---|---|
| Heterozygosity | 1.75375% | 1.80378% |
| Genome Haploid Length | 558,344,824 bp | 565,078,853 bp |
| Genome Repeat Length | 350,461,528 bp | 354,688,339 bp |
| Genome Unique Length | 207,883,296 bp | 210,390,514 bp |
| Model Fit | 85.7458% | 98.2203% |
| Read Error Rate | 0.574939% | 0.574939% |

*Table 4*: Genome metrics for *L. baikalensis,* computed using Genoscope, using Jellyfish-derived *k* mer counts (size = 21 bp).

| ***Eukaryote Cassette:*** | ***Baikalospongia bacillifera* (A2)** | ***Lubomirskia abietina* (A10)** | ***Lubomirskia baikalensis* (A8)** | **SPAdes Assembly** |
|---|---|---|---|---|
| **Complete BUSCOs** | 299 | 297 | 291 | 235 |
| **-Complete (Single Copy)** | 192 | 196 | 178 | 204 |
| **-Complete (Duplicated)** | 107 | 101 | 113 | 31 |
| **Fragmentary BUSCOs** | 2 | 2 | 9 | 51 |
| **Missing BUSCOs** | 2 | 4 | 3 | 17 |
| **Total BUSCO genes** | 303 | 303 | 303 | 303 |
| ***Metazoan Cassette*:** | **A2** | **A10** | **A8** | **SPAdes Assembly** |
| **Complete BUSCOs** | 918 | 914 | 904 | 562 |
| **-Complete (Single Copy)** | 555 | 556 | 515 | 493 |
| **-Complete (Duplicated)** | 363 | 358 | 389 | 69 |
| **Fragmentary BUSCOs** | 19 | 18 | 31 | 247 |
| **Missing BUSCOs** | 41 | 46 | 43 | 169 |
| **Total BUSCO genes** | 978 | 978 | 978 | 978 |

*Table 5:* BUSCO results for genome and transcriptome assemblies.
